# MiDAS 4: A global catalogue of full-length 16S rRNA gene sequences and taxonomy for studies of bacterial communities in wastewater treatment plants

**DOI:** 10.1101/2021.07.06.451231

**Authors:** Morten Simonsen Dueholm, Marta Nierychlo, Kasper Skytte Andersen, Vibeke Rudkjøbing, Simon Knutsson, the MiDAS Global Consortium, Mads Albertsen, Per Halkjær Nielsen

**Affiliations:** Center for Microbial Communities, Department of Chemistry and Bioscience, Aalborg University, Aalborg, Denmark

## Abstract

Biological wastewater treatment and an increased focus on resource recovery is fundamental for environmental protection, human health, and sustainable development. Microbial communities are responsible for these processes, but our knowledge of their diversity and function is still poor, partly due to the lack of good reference databases and comprehensive global studies. Here, we sequenced more than 5 million high-quality, full-length 16S rRNA gene sequences from 740 wastewater treatment plants (WWTPs) across the world and used the sequences to construct MiDAS 4, a full-length amplicon sequence variant resolved 16S rRNA gene reference database with a comprehensive taxonomy from the domain to species-level for all references. Using a study-independent amplicon dataset from the Global Water Microbiome Consortium project (269 WWTPs), we showed that the MiDAS 4 database provides much better coverage for bacteria in WWTPs worldwide compared to commonly applied universal references databases, and greatly improved the rate of genus and species-level classification. Hence, MiDAS 4 provides a unifying taxonomy for the majority of prokaryotic diversity in WWTPs globally, which can be used for linking microbial identities with their functions across studies. Taking advantage of MiDAS 4, we carried out an amplicon-based, global-scale microbial community profiling of activated sludge plants using two common sets of primers targeting the V1-V3 and V4 region of the 16S rRNA gene. We found that the V1-V3 primers were generally best suited for this ecosystem, and revealed how environmental conditions and biogeography shape the activated sludge microbiota. We also identified process-critical taxa (core and conditionally rare or abundant taxa), encompassing 966 genera and 1530 species. These represented approximately 80% and 50% of the accumulated read abundance, respectively, and represent targets for further investigations. Finally, we showed that for well-studied functional guilds, such as nitrifiers or polyphosphate accumulating organisms, the same genera were prevalent worldwide, with only a few abundant species in each genus.

## Introduction

The invention of the activated sludge process for biological treatment of wastewater took place approximately 100 years ago ^1^ and is now the world’s largest application of biotechnology by volume ^2^. The process relies on microbial degradation of organic and inorganic compounds, biotransformation of toxic substances, and removal of pathogens. However, wastewater contains many resources, which are currently poorly exploited. To meet the UN sustainable development goals, a transition is taking place to integrate treatment with the recovery of resources and energy production ^3,4^. Activated sludge and other treatment systems, such as granular sludge and biofilters, all rely on complex microbial communities. Advances in the understanding of the microbial ecology of these microbial communities have been ongoing for decades. However, at the most fundamental level, many important microbes remain unidentified and undescribed.

While several studies have attempted to resolve the microbial diversity in wastewater treatment plants (WWTPs), most have focused on few facilities in specific countries or regions ^5–12^. The only global diversity study of WWTPs concluded that there are billions of different species-level OTUs (97% identity, 16S rRNA V4 region), and that very few OTUs are shared across the world ^2^. The 28 core OTUs identified in the study only accounted for 12% of accumulated read abundance in the samples, suggesting that we deal with an overwhelming microbial diversity and complexity. However, studies of process-critical functional groups have indicated that their global diversity could be much lower, especially if we focus only on the abundant species, which are likely to have a notable impact on treatment performance ^13^.

Nearly all microbial community studies of WWTPs are seriously hampered by several problems that limit our insight and ability to share knowledge: One is the application of different wet-lab protocols, e.g., DNA extraction methods and choice of amplicon primers. This problem can be partly mitigated by using standardized protocols for sampling, DNA extraction, and amplicon library preparation ^14–16^. In the MiDAS project, we have thoroughly evaluated all steps for activated sludge samples and provide detailed protocols online (https://www.midasfieldguide.org/guide/protocols). The second issue is the use of different 16S rRNA gene reference databases that lack reference sequences with high identity to those present in WTTPs for many microbes, and also lack a comprehensive taxonomy for the many uncultured environmental taxa ^17^. To overcome these problems, we recently developed MiDAS 3, an ecosystem-specific full-length 16S rRNA gene reference database for wastewater treatment systems ^17,18^. Although MiDAS 3 is only based on Danish nutrient removal plants and anaerobic digesters, it also performed well on samples from similar plants in other countries^17^. However, more plant designs, process types, and geographical locations are needed to cover the global microbial diversity in WWTPs at the highest taxonomic resolution.

Here we present the largest global WWTP sampling and sequencing campaign to date with samples from 740 WWTPs. More than 5 million high-quality, full-length 16S rRNA gene sequences were obtained and used to expand MiDAS 3 to cover the global diversity of microbes in wastewater treatment systems. The resulting database and taxonomy (MiDAS 4) represent a comprehensive catalogue that may act as a common vocabulary for linking microbial taxonomy with function among studies across the field. Furthermore, we carried out amplicon surveys on all activated sludge samples obtained based on the two commonly applied amplicon primer sets targeting V1-V3 and V4. With this data we (i) evaluate which primer set is generally best suited for microbial community profiling of WWTPs, (ii) determine which environmental and geographic parameters correlate with specific genera, (iii) identify process-important taxa, and (iv) investigate the genus- and species-level diversity within important functional guilds.

## Results and Discussion

The MiDAS global consortium was established in 2018 to coordinate the sampling and collection of metadata from WWTPs across the globe (**Data S1**). Samples were obtained in duplicates from 740 WWTPs in 425 cities, 31 countries on six continents (**Figure 1a**), representing the largest global sampling of WWTPs to date. The majority of the WWTPs were configured with the activated sludge process (69.7%) (**Figure 1b**), and these were the main focus of the subsequent analyses. Nevertheless, WWTPs based on biofilters, moving bed bioreactors (MBBR), membrane bioreactors (MBR), and granular sludge were also sampled to cover the microbial diversity in other types of WWTPs. The activated sludge plants were mainly designed for carbon removal only (C) (22.1%), carbon removal with nitrification (C,N) (9.5%), carbon removal with nitrification and denitrification (C,N,DN) (40.9%), and carbon removal with nitrogen removal and enhanced biological phosphorus removal, EBPR (C,N,DN,P) (21.7%) (**Figure 1c**). The first type represents the most simple design whereas the latter represent the most advanced process type with varying oxic and anoxic stages or compartments.

**Figure 1:**
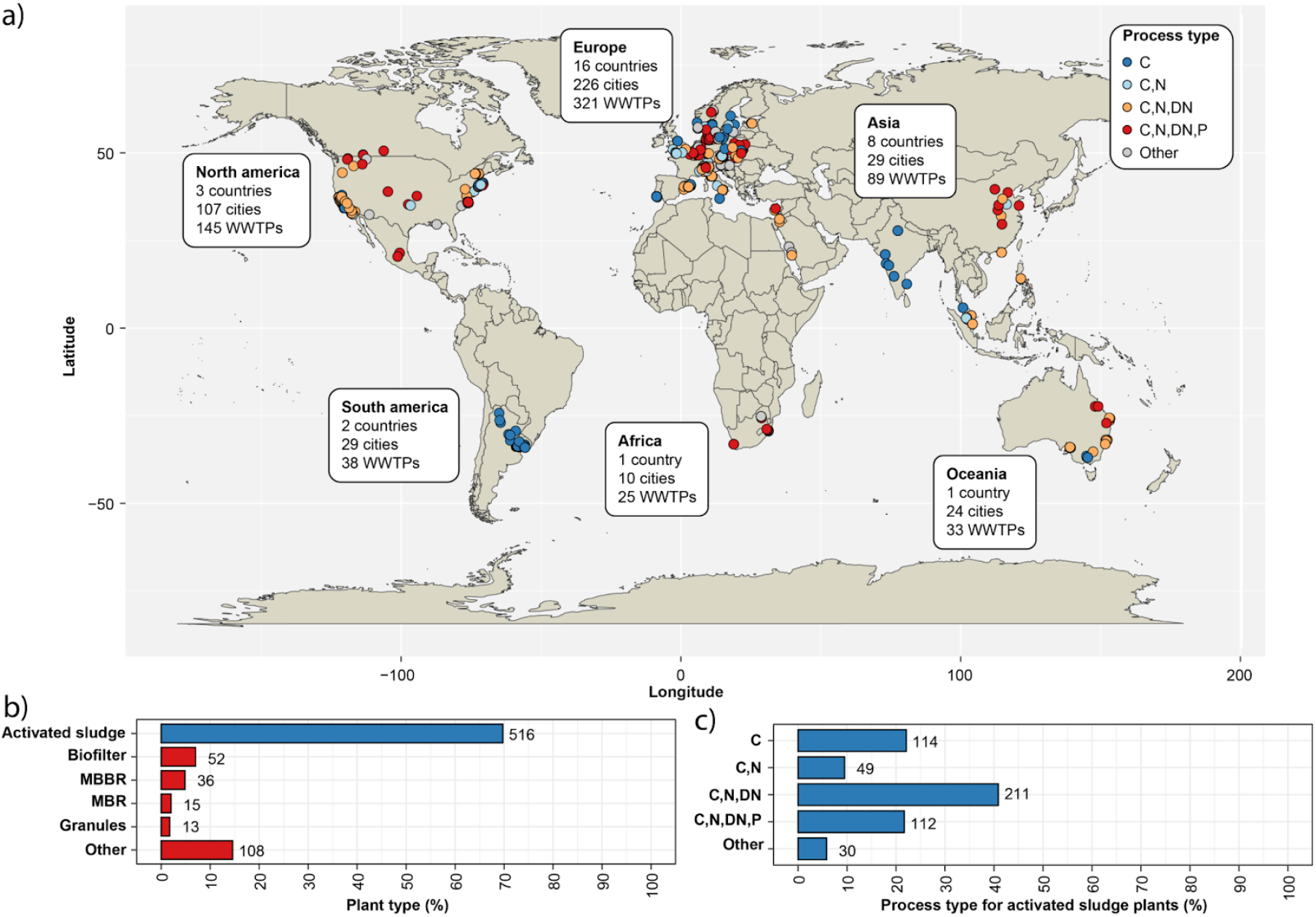
Sampling of WWTPs across the world. a) Geographical distribution of WWTPs included in the study and their process configuration. b) Distribution of WWTP plant types. MBBR: moving bed bioreactor; MBR: membrane bioreactor. c) Distribution of process types for the activated sludge plants. C: carbon removal; C,N: carbon removal with nitrification; C,N,DN: carbon removal with nitrification and denitrification; C,N,DN,P: carbon removal with nitrogen removal and enhanced biological phosphorus removal (EBPR). The values next to the bars are the number of WWTPs in each group.

### MiDAS 4: a new global 16S rRNA gene catalogue and taxonomy for WWTPs

Microbial community profiling at high taxonomic resolution (genus- and species-level) using 16S rRNA gene amplicon sequencing requires a reference database with high-identity reference sequences (≥99% sequence identity) for the majority of the bacteria in the samples and a complete seven-rank taxonomy (domain to species) for all reference sequences ^17,18^. To create such a database for bacteria in WWTPs globally, we applied synthetic long-read full-length 16S rRNA gene sequencing ^17,19^ on samples from all WWTPs included in this study.

More than 5.2 million full-length 16S rRNA gene sequences were obtained after quality filtering and primer trimming. The sequences were processed with AutoTax ^17^, which yielded 80,557 full-length 16S rRNA gene ASVs (FL-ASVs). These were added to our previous MiDAS 3 database ^18^, resulting in a combined database (MiDAS 4) with a total of 90,164 unique FL-ASV reference sequences. Out of these, 88% had best hits in the SILVA 138 SSURef NR99 database above the genus-level threshold (≥94.5% identity) and 56% above the species-level threshold (≥98.7% identity) (**Figure 2b**, **Table 1**).

**Table 1:** Novel sequences and *de novo* taxa observed in the MiDAS 4 reference database. Sequence novelty was determined based on the percent identity between each FL-ASV and their closest relative in the SILVA_138_SSURef_Nr99 database and taxonomic thresholds proposed by Yarza et al.^20^. Taxonomy novelty was defined based on the number of *de novo* taxa assigned by AutoTax at each taxonomic rank.

|  | Sequence novelty |  | Taxonomy novelty* |  |
| --- | --- | --- | --- | --- |
|  | Sequences | Percentage | *De novo* Taxa | Percentage |
| New phylum (<75.0%) | 84 | 0.09% | 26 | 30.59% |
| New class (<78.5%) | 183 | 0.20% | 83 | 37.22% |
| New order (<82.0%) | 334 | 0.37% | 297 | 46.77% |
| New family (<86.5%) | 1,067 | 1.18% | 1,313 | 69.84% |
| New genus (<94.5%) | 10,739 | 11.91% | 8,220 | 86.33% |
| New species (<98.7%) | 40,036 | 44.40% | 30,264 | 96.54% |

\**De novo* species also include known species that cannot be resolved based on full-length 16S rRNA genes.

**Figure 2:**
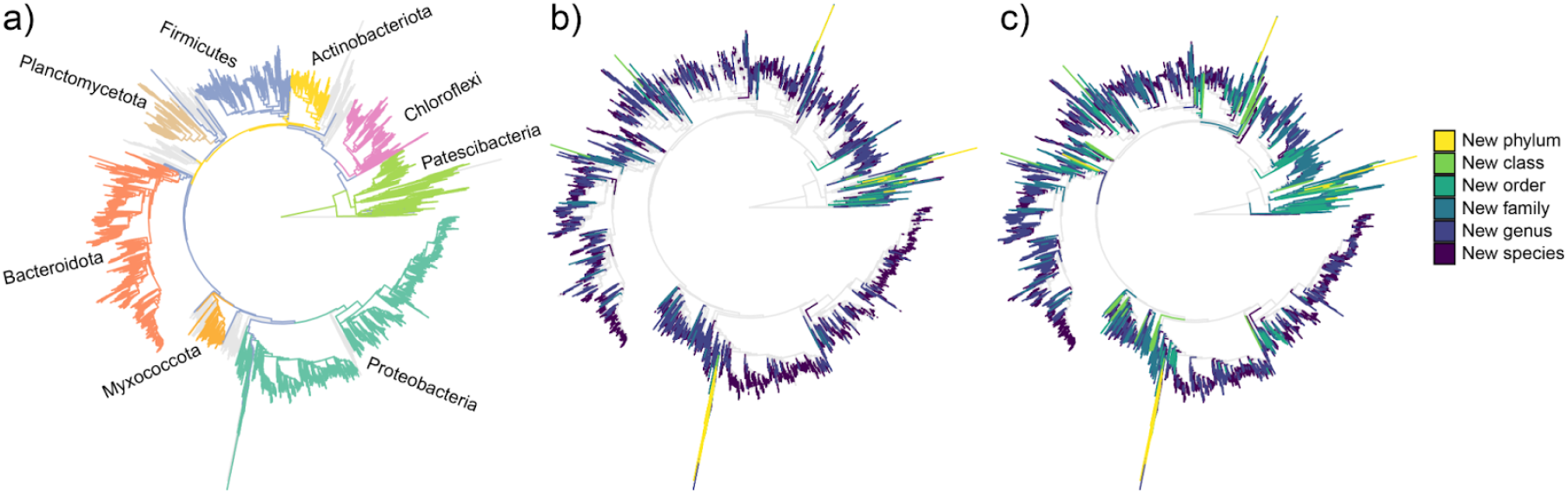
Novel sequences and *de novo* taxa observed in the MiDAS 4 reference database. Phylogenetic trees based on the FL-ASVs in the MiDAS 4 database. a) The eight most diverse phyla, b) Sequence novelty as determined by the percent identity between each FL-ASV and their closest relative in the SILVA_138_SSURef_Nr99 database and taxonomic thresholds proposed by Yarza et al. ^20^. c) Taxonomy novelty defined based on the assignment of *de novo* taxa by AutoTax ^17^.

### MiDAS 4 reveals many novel taxa

Although only a small percentage of the reference sequences in MiDAS 4 represented novel higher rank taxa (phylum, class, or order) according to the sequence identity thresholds proposed by Yarza *et al*. ^20^, a large number of sequences lacked lower-rank taxonomic classifications and was assigned *de novo* placeholder names by AutoTax ^17^ (**Figure 2c, Table 1**). In total, new *de novo* taxonomic names were generated by AutoTax for 26 phyla (30.6% of observed), 83 classes (37.2%), 297 orders (46.8%), and more than 8,000 genera (86.3%).

Phylum-specific phylogenetic trees were created to determine how the FL-ASV reference sequences that were classified as *de novo* phyla were related to previously described taxa (**Figure S1a**). The majority (65 FL-ASVs) created deep branches from within the Alphaproteobacteria together with 16S rRNA gene sequences from mitochondria, suggesting they represented novel mitochondrial 16S rRNA genes rather than true novel phyla. We also observed several FL-ASVs assigned to *de novo* phyla that branched from two classes within the Patescibacteria: the Parcubacteria (3 FL-ASVs) and the Microgenomatis (22 FL-ASVs). These two classes were originally proposed as superphyla due to their unusually high rate of evolution ^21,22^. It is, therefore, likely that the *de novo* phyla are artefacts resulting from the simple approach behind the AutoTax *de novo* taxonomy assignment, which does not take different evolutionary rates into account^17^. Most of the class-, and order-level novelty was found within the Patescibacteria, Proteobacteria, Firmicutes, Planctomycetota, and Verrucomicrobiota. (**Figure S1b**). At the family- and genus-level, we also observed many *de novo* taxa affiliated to Bacteroidota, Bdellovibrionota, and Chloroflexi.

### MiDAS 4 provides a common taxonomy for the field

The performance of the MiDAS 4 database was evaluated based on an independent amplicon dataset from the Global Water Microbiome Consortium (GWMC) project ^2^, which covers approx. 1200 samples from 269 WWTPs. The raw GWMC amplicon data of the 16S rRNA gene V4 region was resolved into ASVs, and the percent identity to their best hits in MiDAS 4 and other reference databases was calculated (**Figure 3a**). The MiDAS 4 database had high-identity hits (≥99% identity) for 72.0%±9.5% (mean±std.dev.) of GWMC ASVs with ≥0.01% relative abundance, compared to 57.9%±8.5% for the SILVA 138 SSURef NR99 database, which was the best of the universal reference databases (**Figure 3a**). Similar analyses of ASVs obtained from the samples included in this study showed, not surprisingly, even better performance with high-identity hits for 90.7%±7.9% of V1-V3 ASVs and 90.0%±6.6% of V4 ASVs with ≥0.01% relative abundance, compared to 60.6±11.9% and 73.9±10.3% for SILVA 138 (**Figure S2a**).

**Figure 3:**
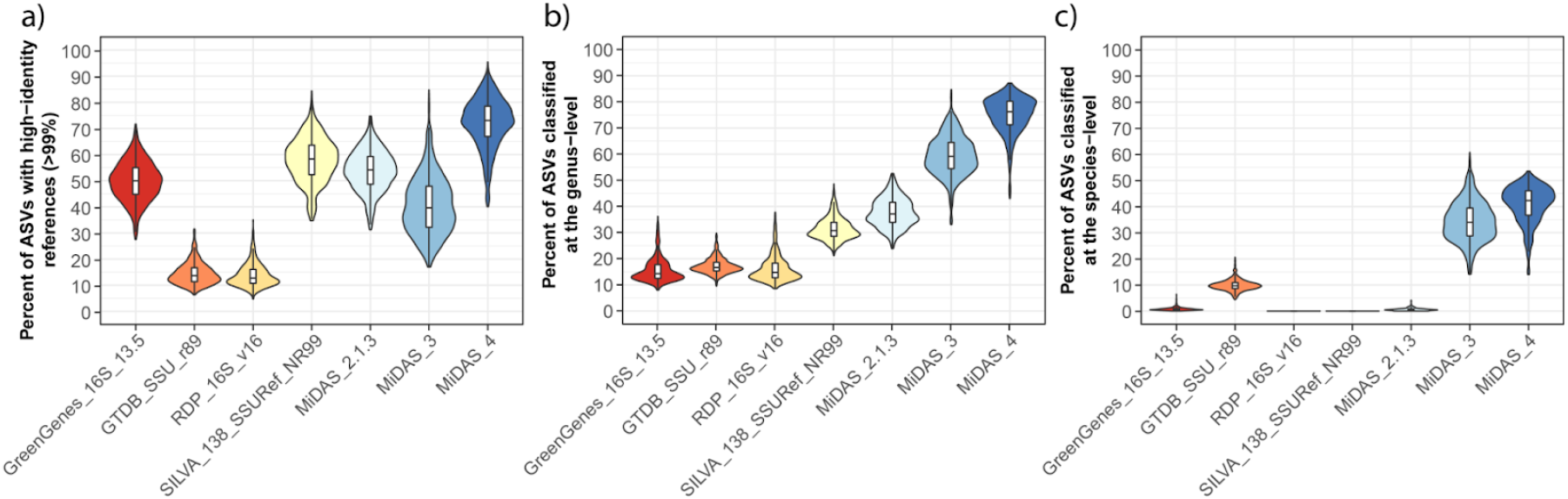
Database evaluation based on amplicon data from the Global Water Microbiome Consortium project. Raw amplicon data from the Global Water Microbiome Consortium project ^2^ was processed to resolve ASVs of the 16S rRNA gene V4 region. The ASVs for each of the 1165 samples were filtered based on their relative abundance (only ASVs with ≥0.01% relative abundance were kept) before the analyses. The percentage of the microbial community represented by the remaining ASVs after the filtering was 88.35% ± 2.98% across samples. a) Percent of ASVs that have high-identity (≥99%) hits in MiDAS 4 and commonly applied universal reference databases. b) and c) Percent of ASVs that received b) genus- or c) species-level classification using sintax with the MiDAS 4 and commonly applied universal reference databases. Outliers have been removed from the boxplots.

Using MiDAS 4 with the sintax classifier, it was possible to obtain genus-level classifications for 75.0%±6.9% of the GWMC ASVs with ≥0.01% relative abundance (**Figure 3b**). In comparison, SILVA 138 SSURef NR99, which was the best of the universal reference databases, could only classify 31.4%±4.2%. When MiDAS 4 was used to classify amplicons from this study, we obtained genus-level classification for 92.0%±4.0% of V1-V3 ASVs and 84.8%±3.6% of V4 ASVs (**Figure S2b**). This is close to the theoretical limit set by the phylogenetic signal provided by each amplicon region analyzed ^17^.

MiDAS 4 was also able to assign species-level classifications to 40.8%±7.1% of the GWMC ASVs. In contrast, GTDB SSU r89, the only universal reference database that contained a comprehensive species-level taxonomy, only classified 9.9%±2.0% of the ASVs (**Figure 3c**). For the ASVs created in this study, MiDAS 4 provided species-level classification for 68.4%±6.1% of the V1-V3 and 48.5%±6.0% of the V4 ASVs (**Figure S2c**).

Based on the large number of WWTPs sampled, their diversity, and the independent evaluation based on the GWMC dataset ^2^, we expect that the MiDAS 4 reference database essentially covers the large majority of bacteria in WWTPs worldwide. Therefore, the MiDAS 4 taxonomy should act as a shared vocabulary for wastewater treatment microbiologists, providing new opportunities for cross-study comparisons and ecological studies at high taxonomic resolution.

### The V1-V3 primer set is recommended for community profiling of WWTPs

Before investigating what factors shape the activated sludge microbiota, we compared short-read amplicon data created for all samples collected in the Global MiDAS project using two commonly used primer sets that target the V1-V3 or V4 variable region of the 16S rRNA gene. The V1-V3 primers were chosen because the corresponding region of the 16S rRNA gene provides the highest taxonomic resolution of common short-read amplicons ^17,23^, and these primers have previously shown great correspondence with metagenomic data and quantitative fluorescence *in situ* hybridization (FISH) results for wastewater treatment systems ^14^. The V4 region has a lower phylogenetic signal, but the primers used to amplify it have better theoretical coverage of the bacterial diversity in the SILVA database ^17,23^.

The majority of genera (62%) showed less than two-fold difference in relative abundances between the two primer sets, and the rest were preferentially detected with either the V1-V3 or the V4 primer (19% for both) (**Figure 4**). However, we observed that several genera of known importance detected in high abundance by V1-V3 were hardly observed by V4, including *Acidovorax, Rhodoferax, Ca.* Villigracilis*, Sphaerotilus*, and *Leptothrix*. In contrast, only a few genera abundant with V4 were strongly underestimated by V1-V3 (e.g., *Acinetobacter* and *Prosthecobacter)*.

**Figure 4:**
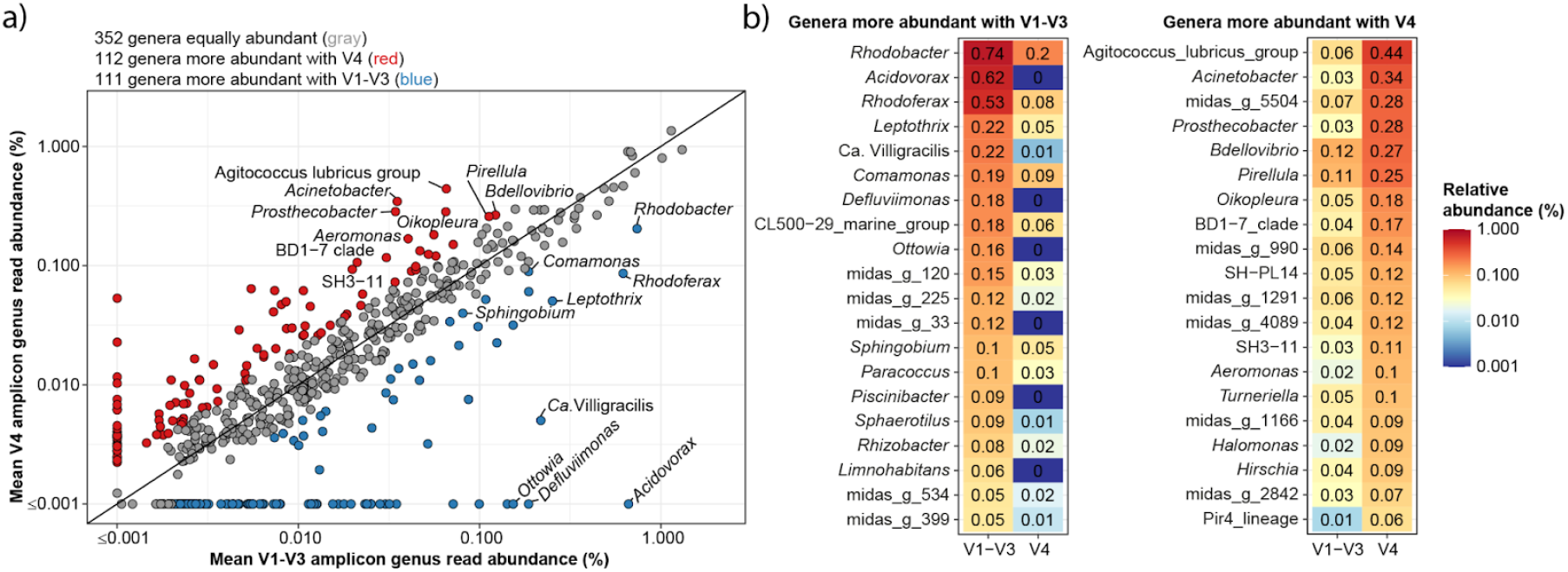
Comparison of relative genus abundance based on V1-V3 and V4 region 16S rRNA gene amplicon data. Mean relative abundance was calculated based on 709 activated sludge samples. a) genera present at ≥0.001% relative abundance in V1-V3 and/or V4 datasets are shown. Genera with less than two-fold difference in relative abundance between the two primer sets are shown with gray circles, and those that are overrepresented by at least two-fold with one of the primer sets are shown in red (V4) and blue (V1-V3). Genus names are shown for all taxa present at minimum 0.1% mean relative abundance (excluding those with *de novo* names). b) Heatmaps of the most abundant genera with more than two-fold relative abundance difference between the two primer sets.

As the V1-V3 primers provide better taxonomic resolution and the least bias in community profiling of abundant process-important taxa, we concluded that this primer set is generally better suited for studies of WWTPs. Accordingly, we primarily focus on the V1-V3 dataset for the following analyses. However, it should be noted that there are cases where the V1-V3 primer set is inappropriate, e.g., for the studies of anammox bacteria ^24,25^. A complete list of differentially detected genera can be found in **Supplementary Data S2**.

### Process and environmental factors affecting the activated sludge microbiota

Alpha diversity analysis revealed that the richness and diversity in activated sludge plants were mainly determined by process type and industrial load (**Figure S3**, **Supplementary results**). The richness and diversity increased with the complexity of the treatment process, as found in other studies, reflecting the increased number of niches ^26^. In contrast, it decreased with high industrial loads, presumably because industrial wastewater often is less complex and therefore promotes the growth of fewer specialised species ^7^.

Distance decay relationship (DDR) analyses were used to determine the effect of geographic distance in the microbial community similarity (**Figure S4**, **Supplementary results**). We found that distance decay was only effective within shorter geographical distances (<2,500 km), which suggests that the microbiota was partly shaped by immigrating bacteria from the source community as recently observed ^27^. In addition, we observed low similarity between geographically separated samples (>2,500 km) at the ASV-level, but higher similarities with OTUs clustered at 97% and even more at the genus-level. This suggests that many ASVs are geographically restricted and functionally redundant in the activated sludge microbiota, so different strains or species from the same genus across the world may provide similar functions.

To gain a deeper understanding of the factors that shape the activated sludge microbiota, we examined the genus-level beta-diversity using principal coordinate analysis (PCoA) and permutational multivariate analysis of variance (PERMANOVA) analyses (**Figure 5**, **Supplementary results**). We found that the overall microbial community was most strongly affected by continent and temperature in the WWTPs. However, process type, industrial load, and the climate zone also had significant impacts. The percentage of total variation explained by each parameter was generally low, indicating that the global WWTPs microbiota represents a continuous distribution rather than distinct states, as observed for the human gut microbiota ^28^.

**Figure 5:**
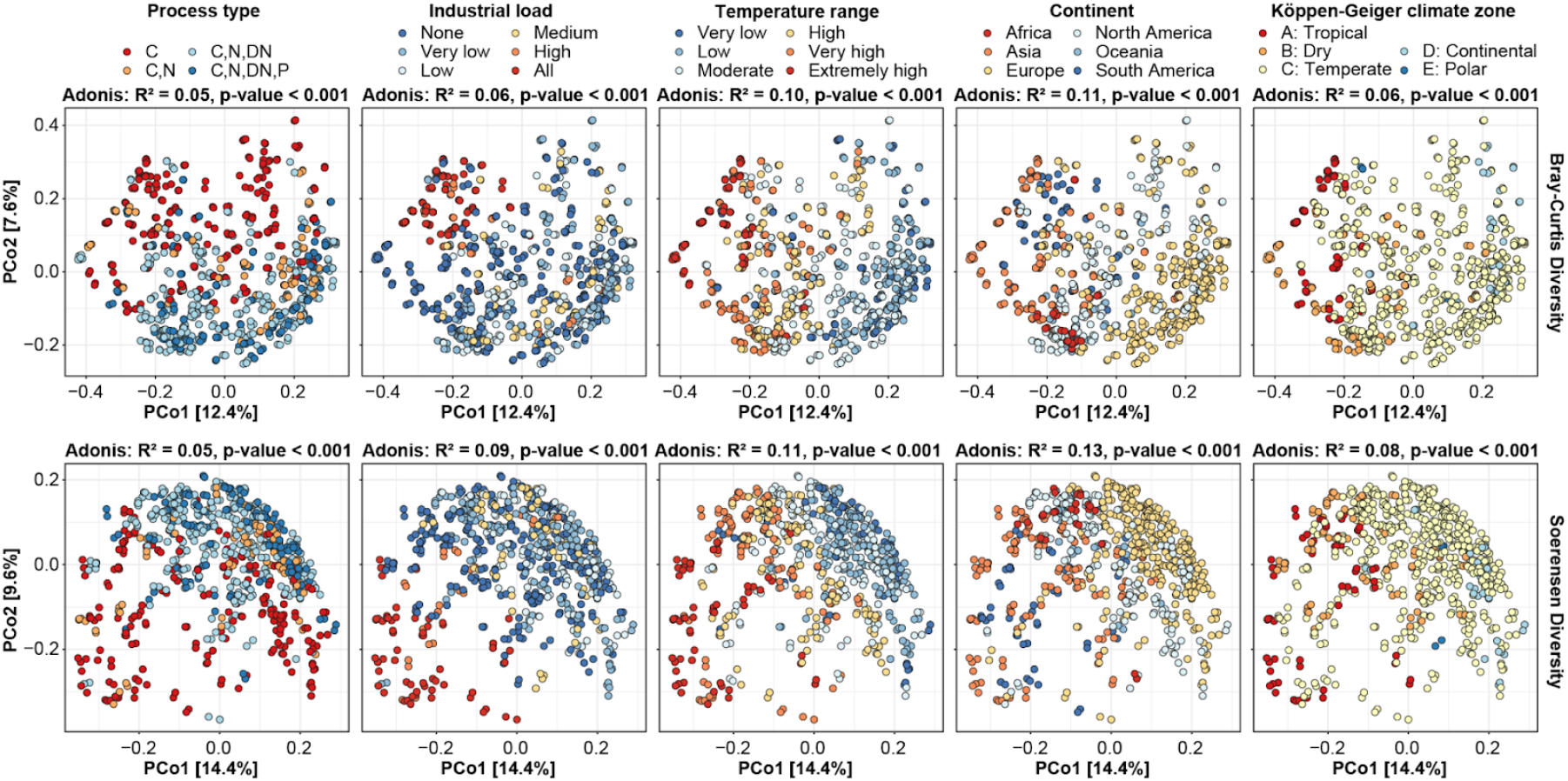
Effects of process and environmental factors on the activated sludge microbial community structure. Principal coordinate analyses of Bray-Curtis and Soerensen beta-diversity for genera based on V1-V3 amplicon data. Samples are colored based on metadata. The fraction of variation in the microbial community explained by each variable was determined by PERMANOVA (Adonis R^2^-values). Temperature range: Very low = <10°C, low = 10-15°C, moderate = 15-20°C, high = 20-25°C, very high = 25-30°C, extremely high = >30°C. Industrial load: None = 0%, very low = 0-10%, low = 10-30%, medium= 30-50%, high 50-100%, all = 100%.

### Genera selected for by process type and temperature

Redundancy analyses (RDA) were used to identify which genera were the strongest indicators for specific processes and/or environmental conditions. RDA was performed on both V1-V3 (**Figure S5**) and V4 (**Figure S6**) amplicon data sets to ensure that essential taxa were not missed due to primer bias. We here highlight the results for process type and temperature. Results for the other parameters and RDA scores for all analyses can be found in **Supplementary Results** and **Supplementary Data S3**, respectively.

The RDA analyses of process types revealed that genera commonly involved in nitrification (*Nitrosomonas* and *Nitrospira*), denitrification (*Rhodoferax*, *Sulfuritalea*), and the polyphosphate accumulating organisms (PAOs) (*Tetrasphaera*, *Ca.* Accumulibacter, and *Dechloromonas*) were strongly enriched in more advanced process types along with *de novo* taxa midas_g_17 (family: Saprospiraceae), midas_g_72 (class: Anaerolineae), and midas_g_300 (order: Sphingobacteriales). Conversely, carbon removal plants were enriched with *Hydrogenophaga* and *Prevotella,* the filamentous genera *Sphaerotilus* and *Thiothrix,* and the glycogen-accumulating organisms (GAOs) *Ca.* Competibacter and *Defluviicoccus*. Specific to the EBPR plants were an increased abundance of known PAOs (see above) and *Azospira*, *Propionivibrio*, *Propioniciclava*, *Ca.* Amarolinea, and the *de novo* taxa midas_g_399 (class: Actinobacteria), midas_g_384 (family: Saprospiraceae), and midas_g_945 (class: Elusimicrobia). The latter genera should be considered targets for further characterization as potential PAOs or GAOs.

The RDA based on temperature showed that high temperatures were associated with an increased abundance of *Ca*. Competibacter, *Thauera*, *Defluviicoccus*, *Azospira*, *Rhodoplanes, Ottowia,* and *Phaeodactylibacter*, whereas lower temperatures favoured the presence of *Flavobacterium*, *Tetrasphaera*, *Ferruginibacter*, *Trichococcus*, *Ca.* Epiflobacter, and *Acinetobacter*. These differences suggest that plants with similar design and operation may have differences in community composition depending on prevailing temperature conditions.

### Core and conditional rare or abundant taxa in the global activated sludge microbiota

Core taxa are commonly defined in complex communities based on how frequent specific taxa are observed in samples from a well-defined habitat ^29^. In addition, an abundance threshold can be applied to select for those taxa that may likely have a quantitative impact on ecosystem functioning ^6^. We here used three frequency thresholds for the core taxa with >0.1% relative abundance in 80% (strict core), 50% (general core), and 20% (loose core) of all activated sludge plants (**Figure 6a**).

**Figure 6:**
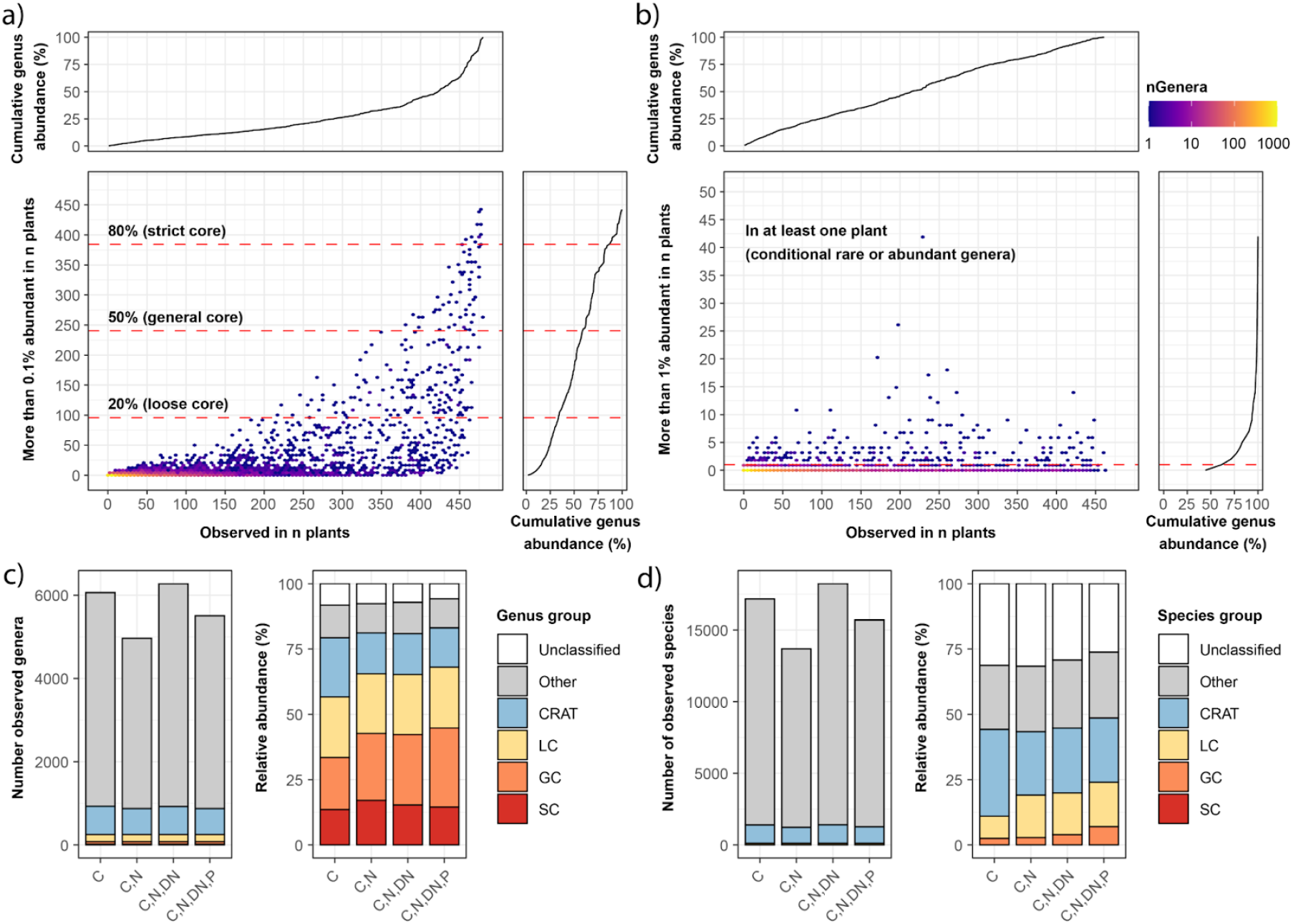
Identification of core and conditionally rare or abundant taxa. a) Identification of strict, general, and loose core genera based on how often a given genus was observed at a relative abundance above 0.1% in WWTPs. b) Identification of conditionally rare or abundant (CRAT) genera based on whether a given genus was observed at a relative abundance above 1% in at least one WWTP. The cumulative genus abundance is based on all ASVs classified at the genus-level. All core genera were removed before identification of the CRAT genera. c) and d) Number of genera and species, respectively, and their abundance in different process types across the global WWTPs. Values for genera and species are divided into strict core (SC), general core (GC), loose core (LC), CRAT, other taxa, and unclassified ASVs.

In addition to the core taxa, we also identified conditionally rare or abundant taxa (CRAT) ^30^ (**Figure 6b**). These are taxa typically present in low abundance, but occasionally become prevalent, including taxa related to process disturbances, such as bacteria causing activated sludge foaming or those associated with the degradation of specific residues in industrial wastewater. CRAT have only been studied in a single wastewater treatment plant treating brewery wastewater, despite their huge potential effect on performance ^30,31^. CRAT are here defined as taxa which are not part of the core, but present in at least one WWTP with a relative abundance above 1%.

Core taxa and CRAT were identified for both the V1-V3 and V4 amplicon data to ensure that critical taxa were not missed due to primer bias. We identified 250 core genera (15 strict, 65 general, and 170 loose) and 715 CRAT genera (**Supplementary Data S4**). The strict core genera (**Figure 7a**) mainly contained genera with versatile metabolisms found in several environments, including *Flavobacterium*, *Novosphingobium*, and *Haliangium*. The general core (**Figure 7b**) included many known bacteria associated with nitrification (*Nitrosomonas* and *Nitrospira*), polyphosphate accumulation (*Tetrasphaera*, *Ca*. Accumulibacter), and glycogen accumulation (*Ca.* Competibacter). The loose core contained well-known filamentous bacteria (*Ca*. Microthrix, *Ca*. Promineofilum, *Ca.* Sarcinithrix, *Gordonia*, *Kouleothrix*, and *Thiothrix*), but also *Nitrotoga,* a less common nitrifier in WWTPs.

**Figure 7:**
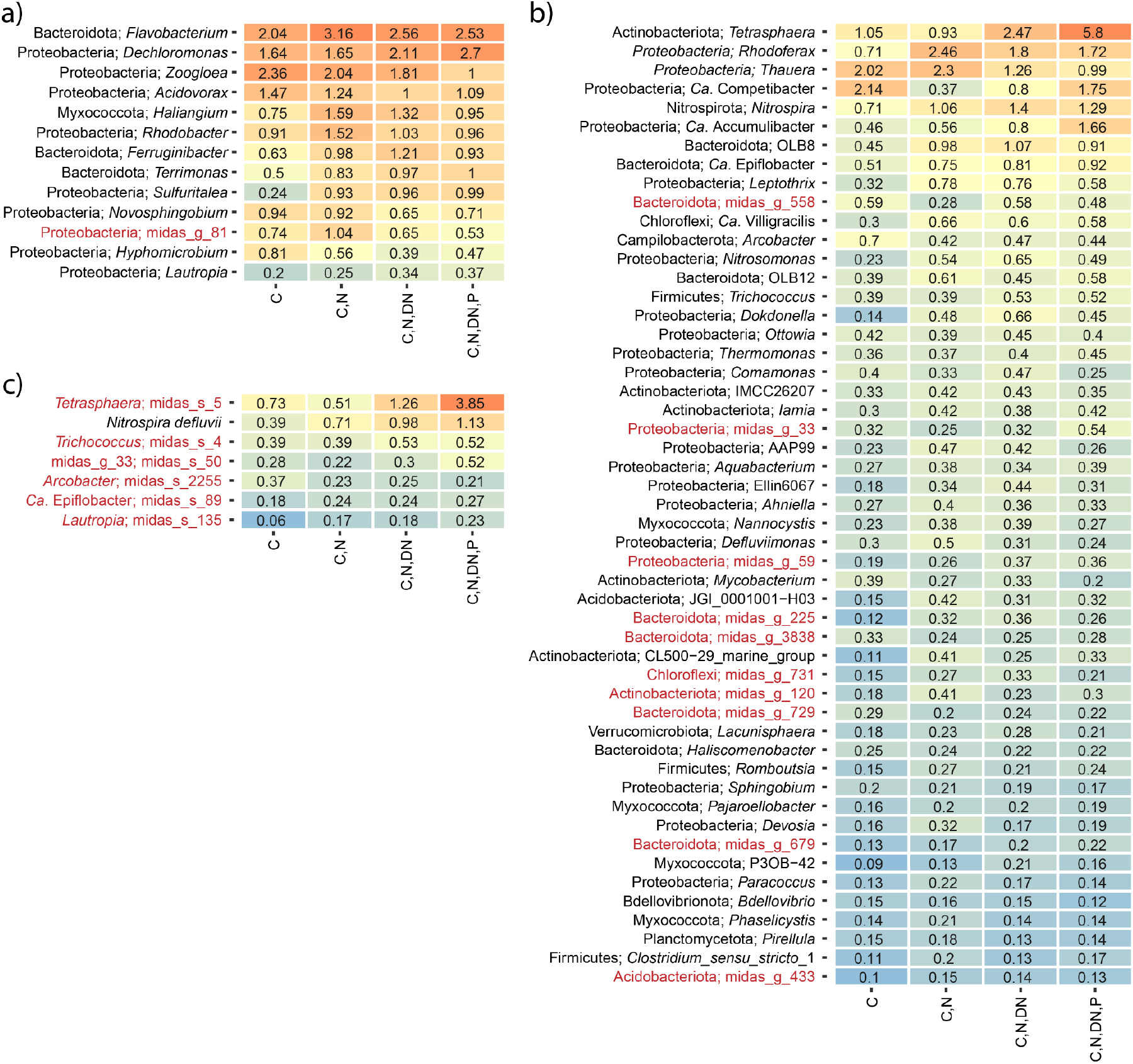
Percent relative abundance of strict and general core taxa across process types. a) Strict core genera. b) General core genera. c) General core species. The taxonomy for the core genera indicates phylum and genus. The genus name is provided together with the core species. *De novo* taxa in the core are highlighted in red.

Because MiDAS 4 allowed for species-level classification, we also identified core and CRAT species based on the same criteria as for genera (**Figure S7, Supplementary Data S4**). This revealed 113 core species (0 strict, 9 general, and 104 loose). The general core species (**Figure 7c**) included the *Nitrospira defluvii* and *Tetrasphaera* midas_s_5, a common nitrifier and PAO, respectively. *Arcobacter* midas_s_2255, a potential pathogen commonly abundant in the influent wastewater, was also part of the general core ^32^. The loose core contained additional species associated with nitrification (*Nitrosomonas* midas_s_139 and *Nitrospira nitrosa*), polyphosphate accumulation (*Ca*. Accumulibacter phosphatis, *Dechloromonas* midas_s_173, *Tetrasphaera* midas_s_45), as well as known filamentous species (*Ca*. Microthrix parvicella and midas_s_2, *Ca*. Villigracilis midas_s_471 and midas_s_9223, *Leptothrix* midas_s_884). In addition to the core species, we identified 1417 CRAT species.

### Many core taxa and CRAT can only be identified with MiDAS 4

The core taxa and CRAT included a large proportion of MiDAS 4 *de novo* taxa. At the genus-level, 106/250 (42%) of the core genera and 500/715 (70%) of the CRAT genera had MiDAS placeholder names. At the species-level, the proportion was even higher. Here placeholder names were assigned to 101/113 (89%) of the core species and 1352/1417 (95%) CRAT species. This highlights the importance of a comprehensive taxonomy that includes the uncultured environmental taxa.

### The core and CRAT taxa cover the majority of the global activated sludge microbiota

Although the core taxa and CRAT represent a small fraction of the total diversity observed in the MiDAS 4 reference database, they accounted for the majority of the observed global activated sludge microbiota (**Figure 6c** and **Figure 6d**). Accumulated read abundance estimates ranged from 57-68% for the core genera and 11-13% for the CRAT, and combined they accounted for 68-79% of total read abundance in the WWTPs depending on process types. The core taxa represented a larger proportion of the activated sludge microbiota for the more advanced process types, which likely reflects the requirement of more versatile bacteria associated with the alternating redox conditions in these types of WWTPs. The remaining fraction, 21-32%, consisted of 6-8% unclassified genera and genera present in very low abundance, presumably with minor importance for the plant performance. The species-level core taxa and CRAT represented 11-24% and 24-33% accumulated read abundance, respectively. Combined, they accounted for almost 50% of the observed microbiota.

### Global diversity within important functional guilds

The general change from simple to advanced WWTPs with nutrient removal and the transition to water resource recovery facilities (WRRFs) requires increased knowledge about the bacteria responsible for the removal and recovery of nutrients, so we examined the global diversity of well-described nitrifiers, denitrifiers, PAOs, and GAOs (**Figure 8**). GAOs were included because they may compete with the PAOs for nutrients and thereby interfere with the biological recovery of phosphorus ^33^. Because MiDAS 4 provided species-level resolution for a large proportion of activated sludge microbiota, we also investigated the species-level diversity within genera affiliated with the functional guilds. A complete overview of species in all genera detected in this global study is provided in the MiDAS field guide (https://www.midasfieldguide.org/guide).

**Figure 8:**
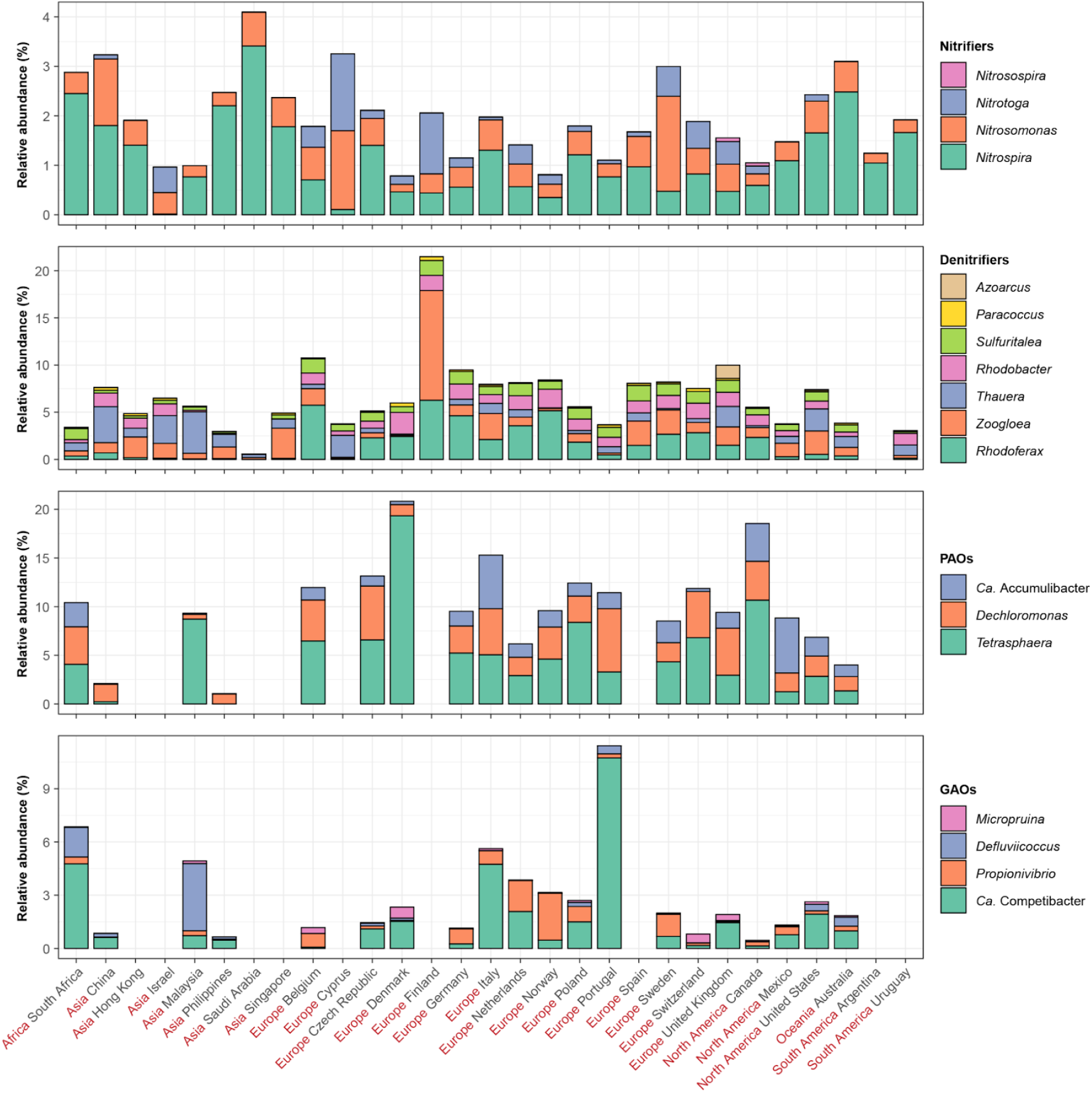
Global diversity of genera belonging to major functional groups. The percent relative abundance represents the mean abundance for each country taking into account only WWTPs with the relevant process types: nitrifiers (C,N; C,ND,N, C,N,DN,P), denitrifiers (C,N,DN, C,N,DN,P), polyphosphate accumulating organisms (PAOs) and glycogen accumulating organisms (GAOs) (C,N,DN,P). The genera are sorted based on their mean global abundance with the most abundant genera at the bottom.

*Nitrosomonas* and potential comammox *Nitrospira* were the only abundant (≥0.1% average relative abundance) genera found among ammonia-oxidizing bacteria (AOB), whereas both *Nitrospira* and *Nitrotoga* were abundant among the nitrite oxidizers (NOB), with *Nitrospira* being the most abundant across all countries (**Figure 8**). *Nitrobacter* was not detected, and *Nitrosospira* was detected in only a few plants in very low abundance (≤0.01% average relative abundance). At the species-level, each genus had 2-5 abundant species (**Figure S8**). The most abundant and widespread *Nitrosomonas* species was midas_s_139. However, midas_s_11707 and midas_s_11733 were dominating in a few countries. For *Nitrospira*, the most abundant species in nearly all countries was *N. defluvii*. ASVs classified as the comammox *N. nitrosa* ^34,35^ was also common in many countries across the world. However, because the comammox trait is not phylogenetically conserved at the 16S rRNA gene level ^34,35^, we cannot conclude that these ASVs represent true comammox bacteria. For *Nitrotoga*, only two species were detected with notable abundance, midas_s_181 and midas_s_9575.

Denitrifying bacteria are very common in advanced activated sludge plants, but are generally poorly described. Among the known genera, *Rhodoferax*, *Zoogloea*, and *Thauera* were most abundant (**Figure 8**). *Zoogloea* and *Thauera* are well-known floc formers, sometimes causing unwanted slime formation ^36^. *Rhodoferax* was the most common denitrifier in Europe, whereas *Thauera* dominated in Asia. The denitrifiers were generally poorly classified at the species-level, except for *Zoogloea* (**Figure S9**). For *Zoogloea*, only *Z. caeni* and midas_s_1080 were abundant.

EBPR is performed by PAOs, with three genera recognized as important in full-scale WWTPs: *Tetrasphaera*, *Dechloromonas*, and *Ca.* Accumulibacter ^13^. According to relative read abundance, all three were found in EBPR plants globally, with *Tetrasphaera* as the most prevalent (**Figure 8**). *Dechloromonas* was also abundant in nitrifying and denitrifying plants without EBPR, indicating a more diverse ecology. Four recognized GAOs were found globally: *Ca.* Competibacter, *Defluviicoccus*, *Propionivibrio,* and *Micropruina*, with *Ca*. Competibacter being the most abundant (**Figure 8**). Only a few species (2-6 species) in each genus were dominant across the world for both PAOs (**Figure S10**) and GAOs (**Figure S11**), except for *Ca*. Competibacter, which covered approx. 20 abundant but country-specific species. Among PAOs, the abundant species were *Tetrasphaera* midas_s_5, *Dechloromonas* midas_s_173, *Ca.* Accumulibacter midas_s_315, *Ca.* A. phosphatis, and *Ca.* A. aalborgensis. Interestingly, some of the most abundant PAOs and GAOs were also abundant in the simple process design with C-removal, indicating more versatile metabolisms.

### Global diversity of filamentous bacteria

Filamentous bacteria are essential for creating strong activated sludge flocs. However, in large numbers, they can also lead to loose flocs and poor settling properties. This is known as bulking, a major operational problem in many WWTPs. Many can also form foam on top of process tanks due to hydrophobic surfaces. Presently, approximately 20 genera are known to contain filamentous species ^37^, and among those, the most abundant are *Ca.* Microthrix, *Leptothrix, Ca*. Villigracilis, *Trichococcus*, and *Sphaerotilus* (**Figure 9**). They are all well-known from studies on mitigation of poor settling properties in WWTPs. Interestingly, *Leptothrix*, *Sphaerotilus* and *Ca.* Villigracilis belong to the genera where abundance-estimation depended strongly on primers, with V4 underestimating their abundance (**Figure 3**). *Ca.* Microthrix and *Leptothrix* were strongly associated with continents, most common in Europe and less in Asia and North America (**Figure 9**).

**Figure 9:**
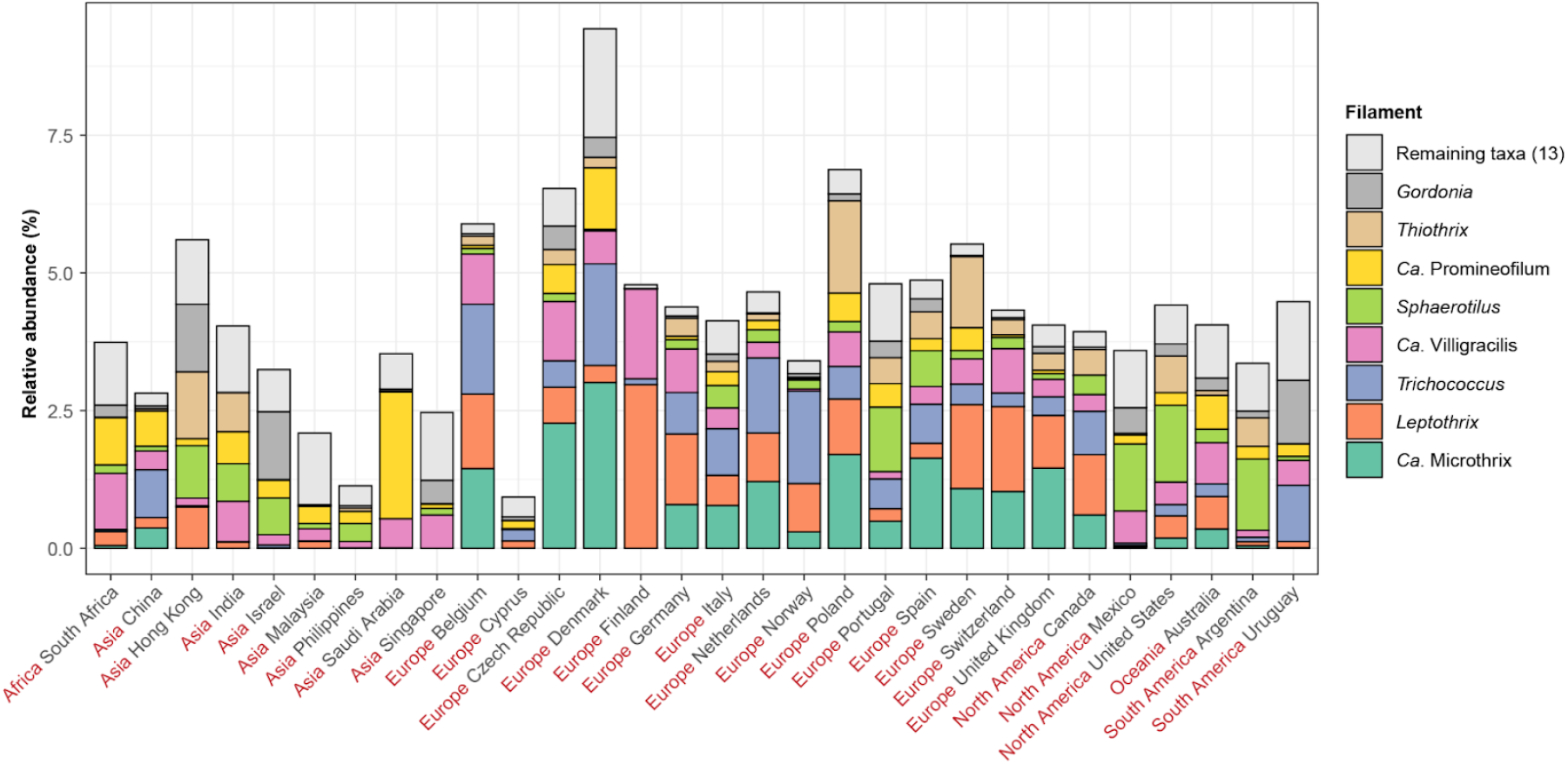
Global diversity of known filamentous organisms. The percent relative abundance represents the mean abundance for each country across all process types. The eight most abundant filamentous organisms based on mean relative abundance across the countries are shown. The filaments are sorted based on their mean global abundance with the most abundant taxa at the bottom. The remaining taxa comprise *Haliscomenobacter*, *Defluviicoccus seviorii*, *Sarcinithrix*, *Ca*. Amarolinea, *Kouleothrix*, *Ca*. Alysiosphaera, *Nocardioides*, midas_g_1668, *Anaerolinea*, *Tetrashaera* midas_s_328, midas_g_105, midas_g_2111, midas_g_344, *Skermania*, *Ca*. Nostocoida, *Neomegalonema*, and *Beggiatoa* (not all were detected).

Many of the filamentous bacteria were linked to specific process types (**Figure S12**), e.g., *Ca*. Microthrix were not observed in WWTPs with carbon removal only, and *Ca*. Amarolinea were only abundant in EBPR plants. The number of abundant species within the genera were generally low, with one species in *Trichococcus,* two in *Ca.* Microthrix and approx. five in *Leptothrix* and *Ca*. Villigracilis (**Figure S13**). Only five abundant species were observed for *Sphaerotilus*. However, a substantial fraction of unclassified ASVs was also observed, demonstrating that certain species within this genus are poorly resolved based on the 16S rRNA gene. *Ca.* Promineofilum was also poorly resolved at the species-level (**Figure S14**).

### Conclusion and perspectives

We present a worldwide collaborative effort to produce MiDAS 4, an ASV-resolved full-length 16S rRNA gene reference database, which covers more than 31,000 species and enables genus- to species-level resolution in microbial community profiling studies. MiDAS 4 covers the vast majority of WWTP bacteria globally and provides a strongly needed common taxonomy for the field, which provides the foundation for comprehensive linking of microbial taxa in the ecosystem with their functional traits.

Presently, hundreds of studies are undertaken to combine engineering and microbial aspects of full-scale WWTPs. However, most ASVs or OTUs in these studies are classified at poor taxonomic resolution (family-level or above) due to the use of incomplete universal reference databases. Because many important functional traits are only conserved at high taxonomic resolution (genus- or species-level), this strongly hampers our ability to transfer new taxa-specific knowledge from one study to another. This will change with MiDAS 4, and we expect that reprocessing of data from earlier studies may reveal new perspectives into wastewater treatment microbiology. Our new online global MiDAS Field Guide presents the data generated in this study and summarises present knowledge about all taxa. We encourage researchers within the field to contribute new knowledge to MiDAS using the contact link in the MiDAS website (https://www.midasfieldguide.org/guide/contact).

The global microbiota of activated sludge plants has been predicted to harbour a massive diversity with up to one billion species ^2^. However, most of these occur at very low abundance and are without importance for the treatment process. By focusing only on the abundant taxa, we can see that the number is much smaller, i.e., approximately 1000 genera and 1500 species. We consider these taxa functionally the most important globally, representing a “most wanted list” for future studies. Some taxa are abundant in most WWTPs (core taxa), and others are occasionally abundant in fewer plants (CRAT). The CRAT have received little attention in the field of wastewater treatment, but they can be of profound importance for WWTP performance. Both groups have a high fraction of poorly characterised species. The new species-level resolution enables us to identify samples where these important core taxa occur in high abundance. This provides an ideal starting point for obtaining high-quality metagenome-assembled genomes (MAGs), isolation of pure cultures, in addition to targeted culture-independent studies to uncover their physiological and ecological roles.

Among the known functional guilds, such as nitrifiers or polyphosphate-accumulating organisms, the same genera were found worldwide, with only a few abundant species in each genus. There were differences in the community structure, and the abundance of dominant species was mainly shaped by process type, temperature, and in some cases, continent. This discovery sends an important message to the field: relatively few species are abundant worldwide, so research or operational results can reliably be transferred from one geographical region to another, stimulating the transition from WWTPs to more sustainable WRRFs.

The relatively low number of uncharacterized abundant species also shows that it is within our reach to describe all of them in terms of identity, physiology, ecology, and dynamics, providing the necessary knowledge for informed process optimization and management. The number of poorly described genera (i.e., those with only a MiDAS placeholder genus name) was 88 among the 250 core genera (35%) and more than 89% at the species-level, so there is still some work to do to link their identities and function. An important step in this direction is the visualization of the populations. With the comprehensive set of FL-ASVs, it is now, for the first time, possible to design comprehensive sets of specific FISH probes, and to critically evaluate the old probes. In the Danish WWTPs, we have successfully done this for groups in the Acidobacteriota ^38^ based on the MiDAS 3 database ^18^. Our recent retrieval of more than 1000 high-quality MAGs from Danish WWTPs with advanced process design is also an important step to link identity to function ^39^. The HQ-MAGs can be linked directly to MiDAS 4 as they contain complete 16S rRNA genes. They cover 62% (156/250) of the core genera and 61% (69/113) of the core species identified in this study. These MAGs may also form the basis for further studies to link identity and function, e.g., by applying metatranscriptomics ^40^ and other *in situ* techniques such as FISH combined with Raman ^41–43^, guided by the “most wanted” list provided in this study. We expect that MiDAS 4 will have huge implications for future microbial ecology studies in wastewater treatment systems.

## Materials and methods

Materials and methods, including statements of data availability and any associated accession codes and references, are available in the supplementary information.

## Acknowledgements

The project has been funded by the Danish Research Council (grant 6111-00617A) and the Villum Foundation (Dark Matter and grant 13351). We thank all the involved WWTPs for providing samples and plant metadata.

## The Global MiDAS consortium

Sonia Arriaga (Environmental Science Department, The Institute for Scientific and Technological Research of San Luis Potosi (IPICYT), San Luis Potosí, Mexico); Rune Bakke (Department of Process, Energy and Environmental Technology, University College of Southeast Norway, Porsgrunn, Norway); Nico Boon (Center for Microbial Ecology and Technology, Ghent University, Ghent, Belgium); Faizal Bux (Institute for Water and Wastewater Technology, Durban University of Technology, Durban, South Africa); Magnus Christensson (Veolia Water Technologies AB, AnoxKaldnes, Lund, Sweden); Adeline Seak May Chua (Department Of Chemical Engineering, Faculty of Engineering, University of Malaya, Kuala Lumpur, Malaysia); Thomas P. Curtis (Environmental Engineering, Newcastle University, Newcastle, England); Eddie Cytryn (The Cytryn Lab, Microbial Agroecology, Volcani Center, Agricultural Research Organization, Rishon Lezion, Israel); Leonardo Erijman (INGEBI-CONICET, Adj. Professor, University of Buenos Aires. Buenos Aires, Argentina); Claudia Etchebehere (Department of Biochemistry and Microbial Genetics, Biological Research Institute “Clemente Estable”, Montevideo, Uruguay); Despo Fatta-Kassinos (NIREAS-International Water Research Center, University of Cyprus, Nicosia, Cyprus); Dominic Frigon (Environmental Engineering, McGill University, Montreal, Canada); Maria Carolina Garcia (School of Microbiology, Universidad de Antioquia, Medellín, Colombia); April Z. Gu (School of Civil and Environmental Engineering, Cornell University, Ithaca, NY, USA); Harald Horn (Water Chemistry and Water Technology and DVGW Research Laboratories, Karlsruhe Institute of Technology (KIT), Karlsruhe, Germany); David Jenkins (David Jenkins & Associates, Inc. Kensington, CA, USA); Norbert Kreuzinger (Institute for Water Quality and Resource Management, TU Wien, Vienna, Austria); Sheena Kumari (Institute for Water and Wastewater Technology, Durban University of Technology, Durban, South Africa); Ana Lanham (Water Innovation and Research Centre, University of Bath, Bath, England); Yingyu Law (Singapore Centre of Environmental Life Sciences Engineering (SCELSE) Nanyang Technological University, Singapore); TorOve Leiknes (Water Desalination and Reuse Center, King Abdullah University of Science and Technology (KAUST), Thuwal, Saudi Arabia); Eberhard Morgenroth (Process Engineering in Urban Water Management, ETH Zürich, Zürich, Switzerland); Adam Muszyñski (Department of Biology, Warsaw University of Technology, Warsaw, Poland); Per H. Nielsen (Department of Chemistry and Bioscience, Aalborg University. Aalborg, Denmark); Steve Petrovski (Environmental Microbial Genetics Lab, La Trobe University, Melbourne, Australia); Maite Pijuan (Technologies and Evaluation Area, Catalan Institute for Water Research, ICRA, Girona, Spain); Suraj Babu Pillai (VA Tech Wabag Ltd., Chennai, India); Maria A.M. Reis (Biochemical engineering group, Universidade Nova de Lisboa, Lisboa, Portugal); Qi Rong (State Key Laboratory of Environmental Aquatic Chemistry, Research Center for Eco-Environmental Sciences, Chinese Academy of Sciences, Beijing, China); Simona Rossetti (Water Research Institute IRSA - National Research Council (CNR), Rome, Italy); Bob Seviour (La Trobe University, Melbourne, Australia); Nick Tooker (Department of Civil and Environmental Engineering, University of Massachusetts Amherst, Amherst, MA, USA); Pirjo Vainio (Kemira Oyj, Espoo R&D Center, Espo, Finland); Mark van Loosdrecht (Environmental Biotechnology, TU Delft, Delft, The Netherlands); R. Vikraman (VA Tech Wabag, Philippines Inc., Makati City, Philippines); Jiøí Wanner (Department of Water Technology and Environmental Engineering, University of Chemistry and Technology, Prague, Czech Republic); David Weissbrodt (Group for Environmental Life Science Engineering, TU Delft. Delft, The Netherlands); Xianghua Wen (School of Environment, Tsinghua University, Beijing, China); Tong Zhang (Environmental Biotechnology Lab Department of Civil Engineering, The University of Hong Kong, Hong Kong).

