## Supplementary information for "MiDAS 4: A global catalogue of full-length 16S rRNA gene sequences and taxonomy for studies of bacterial communities in wastewater treatment plants"

**Affiliation:**

**Running title:** Global microbiota of wastewater treatment plants

**Table of content:**

Page 2-11: Materials and methods
Page 12-14: Supplementary results
Page 15-16: Supplementary references
Page 17-30: Supplementary figure S1-S17

### Materials and methods:

#### *Sampling and metadata collection*

To facilitate sampling of WWTPs across the globe, we established the MiDAS global consortium, which consists of 39 wastewater treatment experts in 33 countries. Members of the consortium acted as national sampling coordinators and were in direct contact with the WWTPs. Two biological sample replicates were obtained from activated sludge aeration tanks or biomass was scraped off biofilters from each WWTP and shipped on ice to the sampling coordinators. For each replicate, 2 mL were preserved in 2 mL RNAlater, stored at 4°C until all national samples were collected (usually within a few days), and then shipped to Aalborg University with cooling elements. Upon arrival, the samples were separated into aliquots that were prepared for nucleic acid purification. Metadata associated with each WWTP was also obtained by the sampling coordinators and is provided as **Data S1**. Minimum information from all plants included continent, country, GPS coordinates, sampling date, temperature in process tank, wastewater composition (municipal vs. industrial COD fraction), process type, and plant type.

#### *General molecular methods*

The concentration and quality of nucleic acids were determined using a Qubit 3.0 fluorometer (Thermo Fisher Scientific) and an Agilent 2200 Tapestation (Agilent Technologies), respectively. Agencourt AMPure XP beads were used as described by the manufacturer, except for the washing steps, where 80% ethanol was used. All commercial kits were used according to the protocols provided by the manufacturer unless otherwise stated.

#### *Nucleic acid purification*

DNA was purified using a custom plate-based extraction protocol based on the FastDNA spin kit for soil (MP Biomedicals). The protocol is available at www.midasfieldguide.org (aau_wwtp_dna_v.8.0). 320 µL of RNAlater preserved sample was pelleted by centrifugation (10,000 x g, 1 min), and the pellet was resuspended in 320 µL PBS and transferred to Lysing Matrix E barcoded tubes (MP Biomedicals). 40 µL MT buffer was added and lysis was performed by bead beating in a FastPrep-96 bead beater (MP Biomedicals) (3x 120 s, 1800 rpm, 2 min incubation on ice between beating). The samples were centrifuged (maximum speed, 10 min) and 200 µL supernatant was transferred to a 96-well PCR-plate. 50 µL Protein Precipitation Solution (PPS) was mixed with each sample, which was then centrifuged again. 150 µL supernatant was cleaned-up using 100 µL Agencourt AMPure XP beads with elution into 60 µL of nuclease-free water. 40 µL of the purified DNA was transferred to a new 96-well plate and stored at -80°C.

#### *Full-length 16S rRNA gene library preparation and sequencing*

Full-length 16S rRNA gene sequencing was carried out using an improved version of our method for synthetic long-read sequencing [1]. Oligonucleotides used can be found in **Table SI-1**. To improve amplification of bacterial 16S rRNA gene diversity for the long-read sequencing, we replaced the 1492r primer previously used [2] with the 1391r primer [3] as it has better coverage of the known bacterial diversity. In addition, we replaced the Taq polymerase used for clonal amplification of the unique molecular identifier (UMI) tagged templates with a proofreading polymerase (see below for *Primary library amplification*). This increased the percentage of error-free reads nearly four-fold, which greatly improves our ability to resolve full-length 16S rRNA ASVs (FL-ASVs) from low abundant taxa because at least two identical sequences are required to resolve an FL-ASV [2] (**Figure SI-1**).

**Table SI-1: Oligonucleotides used for full-length 16S rRNA gene library preparation.** Unique molecular tags, and sample barcodes are marked with blue and red, respectively.

| Name | Sequence | Protocol step |
|---|---|---|
| f16S_pcr1_fw1 | CTCCACCCAGACTCATCCATNNNNNNNNNNNNNNNTGCCTCTTAGAGTTTGATCMTGGCTCAG | Barcoding and unique molecular |
| f16S_pcr1_fw2 | CTCCACCCAGACTCATCCATNNNNNNNNNNNNNNNTCCTCTACAGAGTTTGATCMTGGCTCAG | identifier (UMI) tagging |
| f16S_pcr1_fw3 | CTCCACCCAGACTCATCCATNNNNNNNNNNNNNNNTCATGAGCAGAGTTTGATCMTGGCTCAG | |
| f16S_pcr1_fw4 | CTCCACCCAGACTCATCCATNNNNNNNNNNNNNNNCCTGAGATAGAGTTTGATCMTGGCTCAG | |
| f16S_pcr1_fw5 | CTCCACCCAGACTCATCCATNNNNNNNNNNNNNNNTAGCGAGTAGAGTTTGATCMTGGCTCAG | |
| f16S_pcr1_fw6 | CTCCACCCAGACTCATCCATNNNNNNNNNNNNNNNGTAGCTCCAGAGTTTGATCMTGGCTCAG | |
| f16S_pcr1_rv1 | AGCGCGGCAAAGATGAAGATNNNNNNNNNNNNNNNTGAACCTTGACGGGCGGTGWGTRCA | |
| f16S_pcr1_rv2 | AGCGCGGCAAAGATGAAGATNNNNNNNNNNNNNNNTGCTAAGTGACGGGCGGTGWGTRCA | |
| f16S_pcr2_fw | CTCCACCCAGACTCATCCAT | Library and clonal amplification |
| f16S_pcr2_rv | AGCGCGGCAAAGATGAAGAT | |
| f16S_readtag_fw | CAAGCAGAAGACGGCATACGAGATGTGACTGGAGTTCAGACGTGTGCTCTTCCGATCTCTCCACCCAGACTCATCCAT | Read-tag library preparation |
| f16S_readtag_rv | CAAGCAGAAGACGGCATACGAGATGTGACTGGAGTTCAGACGTGTGCTCTTCCGATCTAGCGCGGCAAAGATGAAGAT | |
| f16S_linktag_fw | CAAGCAGAAGACGGCATACGAGATCGGTCTCGGCATTCCTGCTGAACCGCTCTTCCGATCTCTGAGCCAKGATCAAACTCT | Linked-tag library preparation |
| f16S_linktag_rv | AATGATACGGCGACCACCGAGATCTACACTCTTTCCCTACACGACGCTCTTCCGATCTTGYACWCACCGCCCGTC | |
| f16S_read2_fw | GCTCTTCCGATCTCTCCACCCAGACTCATCCAT | Illumina MiSeq/HiSeq sequencing |
| f16S_read2_rv | GCTCTTCCGATCTAGCGCGGCAAAGATGAAGAT | |

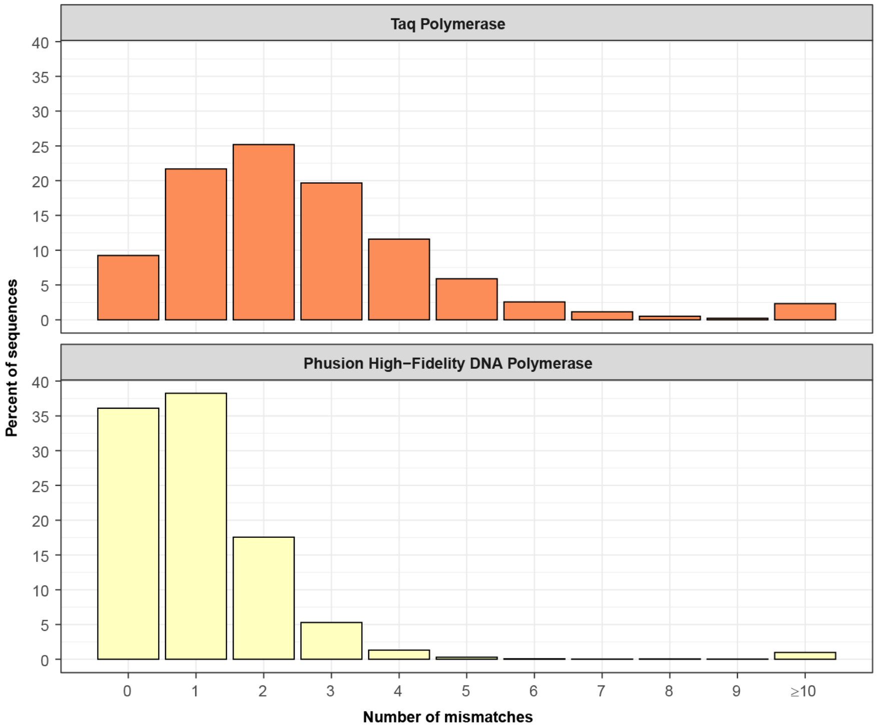


**Figure SI-1: Replacement of the Taq polymerase with the Phusion HiFi DNA polymerase improves the raw synthetic long-read error-rate.** The libraries were created as described in the materials and methods with the exception of the clonal amplification with the Taq polymerase which was carried out as previously described by Karst et al. 2018 [1]. The ZymoBIOMICS Microbial Community DNA Standard was used as a template and sequences were matched against the corrected reference sequences from Karst et al. 2019 [4].

*Adapter annealing by PCR:* Adaptors containing barcodes and UMI and defined primer binding sites were added to each end of the bacterial 16S rRNA genes by PCR. The reaction contained 10 µL of 10x PCR Buffer (Qiagen), 2 µL of 10 mM dNTP (Qiagen), 5 µL of 10 µM f16S_pcr1_fw, 5 µL of 10 µM f16S_pcr1_rv, 4 µL of 25 mM $MgCl_2$, 0.5 µL of 5 U/µL Taq polymerase (Qiagen), 100 ng of pooled template DNA (from two to five WWTPs), and nuclease-free water to 100 µL. The reaction was incubated with an initial denaturation at 94°C for 3 min followed by 2 cycles of denaturation at 94°C for 30 s, annealing at 56°C for 30 s, and extension at 72°C for 3 min, and then a final extension at 72°C for 5 min. The sample was purified using 0.6x AMPure XP beads and eluted in 10 µL nuclease-free water.

*Primary library amplification:* The tagged 16S rRNA gene amplicons were amplified using PCR to obtain enough product for quantification and sequencing. The reaction contained 9 µL of adaptor annealed sample, 20 µL 5x Phusion HF buffer (NEB), 2 µL of 10 mM dNTP, 5 µL of 10 µM f16S_pcr2_fw, 5 µL of 10 µM f16S_pcr2_rv, 4 µL of 25 mM MgCl, 54 µL nuclease-free water, and 1 µL 2U/µL Phusion HF DNA polymerase (NEB). The reaction was incubated with an initial denaturation at 98°C for 30 s followed by 15 cycles of denaturation at 98°C for 10 s, annealing at 62°C for 30 s, and extension at 72°C for 1 min and then a final extension at 72°C for 5 min. The PCR product was purified using 0.6x AMPure XP beads and eluted in 11 µL nuclease-free water. The amplicons were validated on a D5000 screentape and quantified with the Qubit dsDNA HS Assay Kit. Up to 10 samples with different barcodes were pooled with an equal amount of DNA from each sample.

*Clonal library amplification:* Tagged amplicon libraries were diluted to approximately 100,000 molecules/µL and amplified by PCR to obtain clonal copies of each uniquely tagged amplicon molecule. The PCR reaction contained 63.5 µL nuclease-free water, 10 µL 10X PCR buffer (Qiagen), 2 µL 10 mM dNTP, 5 µL 10 µM f16S_pcr2_fw , 5 µL 10 µM f16S_pcr2_rv , 4 µL 25 mM MgCl, 0.5 µL 5 U/µL Taq polymerase (Qiagen) and 10 µL diluted tagged amplicon product. The reaction was initiated by denaturation at 94°C for 3 min, followed by 20 cycles of denaturation at 94°C for 30 s, annealing at 62°C for 30 s and extension at 72°C for 2 min and finishing with final extension at 72°C for 5 min. The PCR product was purified using 0.6x AMPure XP beads with elution into 21 µL nuclease-free water. The product quality and concentration were analysed on a D5000 screentape and with the Qubit dsDNA HS Assay Kit, respectively.

*Read-tag library preparation:* A Nextera library preparation kit (Illumina) was used to prepare a paired-end read-tag sequencing library from the clonal library using a customised protocol. A tagmentation reaction was prepared with 100 ng of the clonal library in 22.5 µL nuclease-free water, 25 µL tagment DNA buffer (Illumina) and 2.5 µL tagment DNA enzyme (Illumina). The reaction was incubated at 55°C for 5 min. The product was immediately diluted to 100 µL and purified using 0.6x AMPure XP beads with elution into 42 µL nuclease-free water.

The tagmentation products were PCR amplified using two separate PCRs (A and B). PCR A selectively amplified fragments containing the 5' termini of the amplicons and PCR B selectively amplified fragments containing the 3' termini. The reactions contained 20 µL purified tagmentation product, 5 µL N504 nextera adaptor (Illumina), 5 µL 10 µM f16S_readtag_fw (PCR A) or f16S_readtag_rv (PCR B) adaptor, 5 µL PCR primer cocktail (Illumina), 10 µL 5x Phusion HF buffer (NEB), 1 µL 10 mM dNTP, 3.5 µL nuclease-free water, and 0.5 µL 2U/µL Phusion HF DNA polymerase (NEB). The following PCR program was used: Initial elongation at 72°C for 3 min, initial denaturation at 98°C for 30 s, and 10 cycles of denaturation at 98°C for 10 s, annealing at

60°C for 30 s and elongation at 72°C for 3 min and finishing with final extension at 72°C for 5 min. The raw read-tag libraries were purified using 1.0x AMPure XP beads with elution into 21 µL nuclease-free water.

To ensure even sequencing coverage across the length of the 16S rRNA gene amplicons, the read-tag libraries were normalised [1]. The libraries were size fractionated on an E-Gel CloneWell gel (Thermo Fisher Scientific). A total of 500 ng GeneRuler 1 kb DNA ladder (Thermo Fisher Scientific) was used as a reference. The gel was run until the 500 bp marker was 1 mm from the elution well, after which 20 µL elution aliquots were sampled and replaced by nuclease-free water every 15 seconds, up to a total of 32 aliquots. Every two aliquots were pooled, yielding 16 pooled aliquots per sample. These were then analysed on an Agilent 2200 Tapestation using the High Sensitivity D1000 Screentape. Fractions with a mean fragment length of 500-1250 bp were used for the pooling. The effective sequencing concentration for fractions from 500-950 bp was determined based on the tapestation data and the empirical formula $C_{seq}$ = Peak molarity [pmol/l] * (-0.0124*(peak size [bp] – 215 bp) + 10.332) [1]. These fractions were pooled in equimolar concentrations ($C_{seq}$). For fractions between 950-1250 bp the entire aliquot was used for pooling (40-50 µL). The pooled aliquots were then purified using 1.0 x AMPure XP beads with elution into 11 µL nuclease-free water. The quality and concentration of the coverage normalised read-tag libraries were analysed on D1000 screentapes and with the Qubit dsDNA HS Assay Kit, respectively.

*Linked-tag library preparation:* Clonal libraries were end-repaired in a reaction containing 20 ng clonal library, 2.5 µL 10x NEBNext End Repair Reaction Buffer (New England Biolabs), 1.25 µL NEBNext End Repair Enzyme Mix (New England Biolabs), and nuclease-free water to 25 µL. The reaction was incubated at 20°C for 30 min. The end-repair reaction was purified using 1.0x AMPure XP beads and eluted into 10 µL nuclease-free water.

The end-repaired sample was circularised in an intra molecular blunt end ligation reaction containing 150 µL nuclease-free water, 20 µL 50% (w/w) PEG 4000 solution (Thermo Fisher Scientific), 20 µL 10x T4 DNA ligase buffer (NEB), 8 µL T4 DNA ligase (NEB), and 2 µL of end-repaired clonal library. The reaction was incubated at 16°C for 60 min. The circularised products were purified using 1.0 x AMPure XP beads and eluted in 10 µL nuclease-free water.

The junction sequence, which contains both unique tags, were amplified by PCR in a reaction containing 8 µL of circularized clonal library, 5 µL 10x PCR buffer (Qiagen), 1 µL 10 mM dNTP mix, 2.5 µL with 10 µM of each f16S_linktag_fw and f16S_linktag_rv,

2 µL 25 mM $MgCl_2$, 30.25 µL nuclease-free water, and 0.25 µL 5u/µL Taq polymerase (Qiagen). The PCR reaction was initiated by denaturation at 94°C for 3 min, followed by 20 cycles of denaturation at 94°C for 20 s, annealing at 56°C for 20 s, and extension at 72°C for 20 s and finishing with a final extension at 72°C for 3 min. The PCR product was purified using 1.0x AMPure XP beads and elution into 12 µL nuclease-free water. The quality and concentration of the linked-tag libraries were analysed on D1000 screentapes and with the Qubit dsDNA HS Assay Kit, respectively.

*Library pooling:* The coverage normalised read-tag library A and B were diluted to 0.9 ng/µL. The linked-tag library was diluted to 0.2 ng/µL. The libraries were pooled by combining 4.6 µL read-tag library A, 4.6 µL read-tag library B, and 0.8 µL linked-tag library.

*Sequencing:* The libraries were paired-end (1 x 240 bp and 1 x 25 bp) sequenced on a HiSeq 2500 instrument (Illumina) using on-board clustering and rapid run mode with a HiSeq PE Rapid Cluster Kit v2 (Illumina) and HiSeq Rapid SBS Kit v2, 200 cycles (Illumina). The SBS reagents were supplemented with 9.5 mL Incorporation Master Mix, 9.5 mL Cleavage Reagent Mix and 7 mL Universal Scan Mix to enable sequencing of 265 cycles. The HiSeq was running HiSeq Control Software v2.2.68 (Illumina) and Real Time analysis v1.18.66.3 (Illumina). The libraries were prepared and loaded on the HiSeq using the standard procedures (Illumina: manual # 15035786 v01; manual # 15050107 v02; manual # 15061846 v01) with the following changes. A volume of 10 µL library pool was denatured by adding 10 µL 0.1 N NaOH solution, mixing well by pipetting and incubating for 5 min at 25°C. The denatured library pool was diluted by adding 980 µL of cold Hybridization Buffer (Illumina). 400 µL of the denatured and diluted library pool was mixed with 20 µL of denatured and diluted 10 pM PhiX control v3 library (Illumina), and stored on ice until loading. Custom read2 primer mix was prepared by mixing 25 µL of 100 µM f16S_read2_fw and 25 µL of 100 µM f16S_read2_rv in a conical tube (15 mL) and diluted with 4,950 µL Hybridization Buffer (final concentration 0.5 µM). When the paired-end reagent rack was loaded on the HiSeq, the Illumina primer mix in position nr. 16 was replaced with the custom read2 primer mix prepared above. When setting up the HiSeq run in the control software, the standard procedure was followed except for the following steps: For the "Recipe Screen", the following options were chosen: Index type options = No Index. Read 1 cycles = 240. Read 2 cycles = 25. After sequencing, bcl2fastq v2.17.1.14 (Illumina) was used to generate fastq files from bcl files using standard settings (manual # 15038058 RevB).

***Short-read amplicon sequencing***

V1-V3 amplicons were made using the 27F (5'-AGAGTTTGATCCTGGCTCAG-3')[5] and 534R (5'-ATTACCGCGGCTGCTGG-3')[7] primers with barcodes and Illumina adaptors (IDT) [6]. 25 µL PCR reactions in duplicate were run for each sample using 1X PCRBIO Ultra Mix (PCR Biosystems), 400 nM of both forward and reverse primer, and 10 ng template DNA. PCR conditions were 95°C, for 2 min followed by 20 cycles of 95°C for 20 s, 56°C for 30 s, and 72°C for 60 s, followed by a final elongation at 72°C for 5 min. PCR products were purified using 0.8 x AMPure XP beads and eluted in 25 µL nuclease-free water.

V4 amplicons were made using the 515F (5'-GTGYCAGCMGCCGCGGTAA-3')[7] and 806R (5'-GGACTACNVGGGTWTCTAAT-3')[8] primers. 25 µL PCR reactions in duplicate were run for each sample using 1X PCRBIO Ultra Mix (PCR Biosystems), 400 nM of both forward and reverse primer, and 10 ng template DNA. PCR conditions were 95°C, for 2 min followed by 30 cycles of 95°C for 15 s, 55°C for 15 s, and 72°C for 50 s, followed by a final elongation at 72°C for 5 min. PCR products were purified using 0.8x AMPure XP beads and eluted in 25 µL nuclease-free water. 2 µL of purified PCR product from above was used as template for a 25 µL Illumina barcoding PCR reaction containing 1X PCRBIO Reaction buffer, 1 U PCRBIO HiFi Polymerase (PCR Biosystems) and 10 µL of Nextera adaptor mix (Illumina). PCR conditions were 95°C, for 2 min, 8 cycles of 95°C for 20 s, 55°C for 30 s, and 72°C for 60 s, followed by a final elongation at 72°C for 5 min. PCR products were purified using 0.8x AMPure XP beads and eluted in 25 µL nuclease-free water.

16S rRNA gene V1-V3 and V4 amplicon libraries were pooled separately in equimolar concentrations and diluted to 4 nM. The amplicon libraries were paired-end sequenced (2 × 300 bp) on the Illumina MiSeq using v3 chemistry (Illumina, USA). 10–20% PhiX control library was added to mitigate low diversity library effects.

***General bioinformatic methods***

Usearch v.11.0.667 [9] was used for processing of 16S rRNA gene amplicon data and for read mapping. Mapping of sequences to references was done with the -usearch_global command and the -id 0, -maxaccepts 0, -maxrejects 0, -top_hit_only, and -strand plus options unless otherwise stated. UNOISE3 [10] was used to resolve exact amplicon sequence variants (ASVs). SINTAX [11] was used for classification of ASVs using the usearch -sintax command with the -strand plus and -sintax_cutoff 0.8 options. Data were analyzed with R v.3.6.3 [12] through RStudio IDE [13], with the tidyverse v.1.2.1 (https://www.tidyverse.org/), vegan v. 2.5 [14], and Ampvis2 v.2.4.0 [15] packages.

***Assembly of full-length 16S rRNA genes***

Raw sequence reads were binned, based on the unique molecular identifiers, *de novo* assembled into the synthetic long-read rRNA gene sequences as previously described [2]. The assembled 16S rRNA gene sequences were oriented based on the SILVA 138 SSURef Nr99 database using the usearch v.11.0.667 -orient command, and trimmed between the 27f and 1391r primer binding sites using the trimming function in CLC genomics workbench v. 20.0. Sequences where both the primer bindings sites were not found were discarded.

***Generation of the MiDAS 4 16S rRNA gene reference database***

The trimmed full-length 16S rRNA genes from above were processed with AutoTax v. 1.5.2 [3] to create full-length 16S rRNA gene amplicon sequence variants (FL-ASV) and these were added to the MiDAS 3 reference database [16] to create MiDAS 4.

***Processing of short-read amplicon data***

16S rRNA gene V1-V3 forward and reverse reads were merged using the usearch -fastq_mergepairs command, filtered to remove phiX sequences using usearch -filter_phix, and quality filtered using usearch -fastq_filter with -fastq_maxee 1.0. Dereplication was performed using -fastx_uniques with -sizeout, and amplicon sequence variants (ASVs) were resolved using the usearch -unoise3 command. OTUs clustered at 97% identity were created using the -cluster_otus command. ASV- and OTU-tables were created by mapping the quality filtered reads to the ASVs using the usearch -otutab command with the -otus or -zotus and -strand plus options. Taxonomy was assigned to OTUs and ASVs using MiDAS 4.8 and the usearch -sintax command with -strand both and -sintax_cutoff 0.8 options.

16S rRNA gene V4 forward reads (reverse reads in relation the 16S rRNA gene) were trimmed with cutadapt v. 2.8 [17] based on the V4 primers with the -g ^GGACTACHVGGGTWTCTAAT...TTACCGCGGCKGCTGGCAC and --discard-untrimmed options. The trimmed reads, which span the entire V4 amplicon, were reverse complemented with usearch -fastx_revcomp, and quality filtered using usearch -fastq_filter with -fastq_maxee 1.0. Subsequent processing was similar to that for the V1-V3 amplicons.

Raw V4 amplicon data from the Global Water Microbiome Consortium [18] was downloaded from NCBI Sequence Read Archive (SRA) with accession number PRJNA509305 using the SRA-Toolkit v. 2.9.2. Forward and reverse reads were merged using the usearch -fastq_mergepairs command, filtered for phiX sequences using usearch

-filter_phix, and quality and adapter filtered using usearch -fastq_filter with the -fastq_stripleft 21, -fastx_stripright 25, and -fastq_maxee 1.0 options. Subsequent processing was similar to that for the V1-V3 amplicons.

***Construction of phylogenetic trees***

FL-ASVs aligned to the global SILVA 138 NR99 alignment were obtained from the AutoTax output (temp/FL-ASVs_SILVA_aligned.fa) and loaded into the SILVA 138 NR99 ARB-database. All bacterial FL-ASVs were selected and exported as a FASTA-alignment using the ssuref:bacteria positional variability by parsimony filter. A tree was created from the alignment using RAxML v. 8.2.12 with the raxmlHPC -PTHREADS -m GTRCAT -n score-f -F -p 32323 command and loaded into ARB.

***Microbial community analyses***

Microbial community analyses were performed for samples collected by the MiDAS global consortium. Only activated sludge plants, which include conventional activated sludge (CAS) and sequence batch reactors (SBR), were chosen for detailed analyses. We further selected plants designed for carbon removal (C), carbon removal with nitrification (C,N), carbon removal with nitrification and denitrification (C,N,DN), and carbon removal with nitrogen removal and enhanced biological phosphorus removal, EBPR (C,N,DN,P). Samples with less than 10,000 reads were discarded, providing in total 861 V1-V3 samples and 666 V4 samples.

Associations between the activated sludge microbiota and the following process-related or environmental variables were investigated: process type (as listed above); industrial load (expressed as fraction of the influent COD); temperature in the process tank (°C), continent and Köppen–Geiger climate classification [19]. Industrial load and temperature was treated as a discrete variable with the following ranges applied: very low (1.8-10.0°C), low (10.1-15.0°C), moderate (15.1-20.0°C), high (20.1-25.0°C), very high (25.1-30.0°C), extremely high (30.1-38.0°C) for temperature; and none (0%), very low (1-10%), low (11-29%), medium (30-50%), high (51-99%), all (100%) for industrial load. The following climate zone groups were used in the analyses: A: tropical/megathermal climates, B: dry (desert and semi-arid) climates, C: temperate/mesothermal climates, D: continental/microthermal climates, E: polar climates.

For alpha-diversity analyses, samples were rarefied to 10,000 reads, and alpha-diversity (Observed taxa and inverse Simpsons) was calculated using the ampvis2 package [15]. The Kruskal-Wallis with Dunn's post-hoc test (Bonferroni correction with α=0.01 before correction) was used to determine statistically significant differences in alpha-diversity between samples grouped by process and environmental variables.

Distance decay relationship was determined using untransformed values of geographic distance against microbial community similarity distance (Bray-Curtis, or Soerensen) for ASVs, 97% OTUs, and genera. Geographical distances between samples were calculated using the distm function in the geosphere R package [20] using Haversine formula. To examine the strength of correlation between geographic and community distance matrices, the Mantel test using Spearman correlation and 999 permutations was performed using the mantel function in the vegan R package.

Beta-diversity distances based on Bray-Curtis (abundance-based) and Soerensen (occurrence-based) for genera (relative genus abundance >0.01% for Soerensen diversity) was calculated using the vegdist function in the vegan R package [14] and visualized by PCoA and RDA plots with the ampvis2 [15] package. Individual process or environmental variables were used as constraints for the RDA. To determine how much individual parameters affected the structure of the microbial community across the WWTPs, a permutational multivariate analysis of variance (PERMANOVA) test was performed on the beta-diversity matrices using the adonis function in the vegan package with 999 permutations.

Core genera and species were defined based on their relative abundances in individual WWTPs. Taxa were defined as abundant when present at >0.1% relative abundance in individual WWTPs. Based on how frequently taxa were observed to be abundant, we defined the following core communities: *loose core* (>20% of WWTPs), *general core* (>50% of WWTPs), and *strict core* (>80% of WWTPs). Additionally, we defined *conditionally rare or abundant taxa (CRAT)* [21] composed of taxa present in one or more WWTPs at >1% relative abundance, but not belonging to the *core taxa*.

***Data and code availability***

Raw and assembled sequencing data is available at the European Nucleotide Archive (https://www.ebi.ac.uk/ena) under the project number (to be published after peer-review, reviews are welcome to get access). The MiDAS 4 reference database in SINTAX and QIIME format, is available at https://www.midasfieldguide.org/guide/downloads. R-markdown scripts used for data analyses and figures are available at https://github.com/msdueholm/Publications/tree/master/Dueholm2021a.

**Supplementary Results:**

***Detailed analysis of alpha-diversity***

Alpha-diversity analyses revealed that the richness and diversity of WWTPs were mainly determined by the process type, industrial load, and continent, whereas the temperature range and climate zone were less discriminatory (**Figure S5**). Both the richness and the diversity increased with the complexity of the treatment process as also found in other studies, reflecting the increased number of niches [22]. In contrast, it decreased with high industrial loads, reflecting that industrial wastewater often is less complex and therefore promotes growth of fewer species [23]. The highest richness and diversity was observed in Oceania and Africa, and the lowest in South America. These differences may partially be linked to differences in process type and industrial load, as the sampled WWTPs in Oceania and Africa were mainly advanced plants (C,N,DN or C,N,DN,P), receiving only little industrial wastewater. In contrast, the South American plants were simple carbon-removal plants that treated >50% industrial wastewater on average.

***Distance decay relationship***

The similarity between microbial communities is generally considered to decrease with geographic distance for most environments including WWTPs [18,24–26]. Therefore, we performed distance decay relationship (DDR) analyses to determine how it affects the activated sludge microbiota on a global scale (**Figure S6**). We observed similar DDR for both ASVs and OTUs (clustered at 97% identity) based on Mantel R values. In addition, we observed that distance decay was only effective within shorter geographical distances (<2,500 km), and almost absent at global scales (>2,500 km).

Although ASVs and OTUs displayed similar DDR, we observed that the Bray-Curtis and Soerensen similarities were considerably lower for ASVs than for OTUs. A complete lack of similarity was observed between many samples at the ASV resolution (similarity $\approx 0$) but not for OTUs. This lack of similarity was especially pronounced for the abundance-weighted Bray-Curtis similarity, indicating that the abundant process-critical bacteria were especially geographically restricted at the ASV level (**Figure S6**).

Because of the low similarity between microbial communities at the ASV level and because OTUs are hard to compare across studies, we also investigated the effect of geographic distance on the taxonomic diversity at the genus level, where we expect that many important traits are conserved (**Figure S6**). This analysis was only possible because of the high classification rate achieved with MiDAS4. The genus level diversity was less affected by DDR compared to ASV and OTU diversity, according to lower Mantel R values. Furthermore, the genus level similarity between the global samples was also markedly higher than the ASV and OTU level diversity.

***Detailed analysis of genus-level beta-diversity***

To gain a deeper understanding of the factors that shape the activated sludge microbiota, we examined the Bray-Curtis (weighted) and Soerensen (unweighted) beta-diversity at the genus-level using principal coordinate (PCoA) and permutational multivariate analysis of variance (PERMANOVA) analyses (**Figure 5**). The PCoA analyses revealed clear separation of the microbial communities in respect to all parameters investigated (**Figure 5**). In addition, we noted clear associations between many of the process-specific and environmental factors investigated, e.g., between high industrial load and simpler process types, and not unexpectedly, between temperature range and climate zones. As a result of these associations, it is hard to accurately assign the contribution of variance by individual factors to the overall beta-diversity. However, by examining how much of the total variance could be explained by each parameter isolated, it was possible to rank the strength of each parameter. We found that the overall microbial community was most strongly affected by continent and temperature in the WWTPs. However, process type, industrial load, and the climate zone also had significant impacts. The percentage of total variation explained by each individual parameter was generally low, indicating that the global WWTPs microbiota represents a continuous distribution rather than distinct states, as has also been observed for the human gut microbiota [27].

***Redundancy analysis for industrial load, continents, and climate zones***

In order to identify which genera were most affected by process and environmental factors, we performed redundancy analyses (RDA) with the relative abundance of each genus as dependent variables and the process and environmental factors as explanatory variables. The RDAs were constrained by each factor separately, and both V1-V3 (**Figure S7**) and V4 (**Figure S8**) amplicon data were analyzed to ensure that important taxa were not missed due to primer bias. RDA scores for all genera and analyses can be found in **Supplementary Data S3**.

When industrial WWTPs (high industrial load) were compared to municipal plants (medium load or below), a reduced abundance of genera associated with advanced process types were observed in plants with high industrial loads (high or all), and an increased abundance of *Defluviicoccus*, *Hyphomicrobium*, *Rhodoplanes*, *Planctomicrobium*, *Ca.* Alysiosphaera, *Azospirillum*, *Azovibrio*, and *Pedomicrobium* were observed. *Defluviicoccus* was also enriched in carbon removal only plants, which is the main process type for treatment of industrial wastewater. *Hyphomicrobium* frequently occurs in WWTPs containing or supplemented with methanol due to its ability to grow on one-carbon compounds [28].

Comparison of continents revealed clear similarities between South America and Asia, and to a lesser degree between Europe and North America. Genera with increased relative abundance in South America and Asia included *Defluviicoccus*, *Ca.* Competibacter, *Thauera*, *Rhodoplanes*, *Phaeodactylibacter,* and *Chitinivorax*, whereas WWTPs in Europe and North America displayed increased abundance of *Rhodoferax*, *Flavobacterium*, *Ca.* Microthrix, *Tetrasphaera,* and *Acidovorax*.

Climate zones showed a clear separation between tropical, dry, and continental climates, whereas the temperate climate had similarities with all the others. Tropical and dry climates were characterized by an increase of abundance of *Ca.* Competibacter, *Defluviicoccus, Thauera*, *Rhodoplanes*, *Ottowia*, UTCFX1, and the *de novo* taxa midas_g_70, midas_g_9648, and midas_g_399. *Rhodoplanes* was the only genus which was more abundant in the tropical climate. Continental climates were characterized by an increased abundance of *Flavobacterium*, *Rhodoferax*, *Rhodobacter, Ferruginibacter, Acidovorax*, *Tetrasphaera*, and the filamentous *Leptothrix* and *Ca*. Microthrix.

### Supplementary Figures

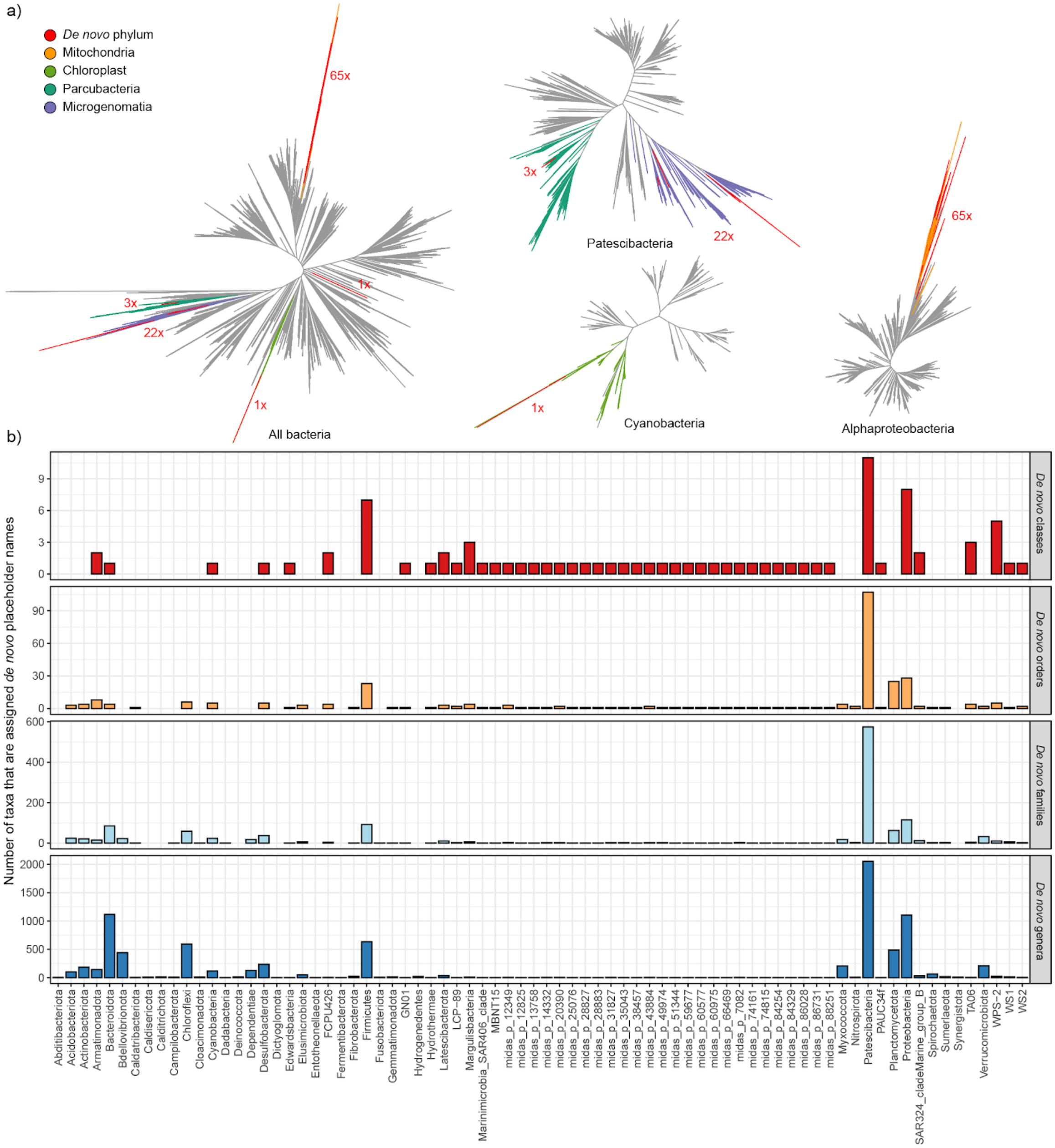


**Figure S1: Phylogenetic and taxonomic location of *de novo* taxa**. a) Phylogenetic trees showing the proposed branching point for FL-ASVs associated with *de novo* phyla. The numbers indicate how many unique FL-ASVs are found at the given location. b) The number of taxa which obtained placeholder names from the MiDAS 4 *de novo* taxonomy in different phyla.

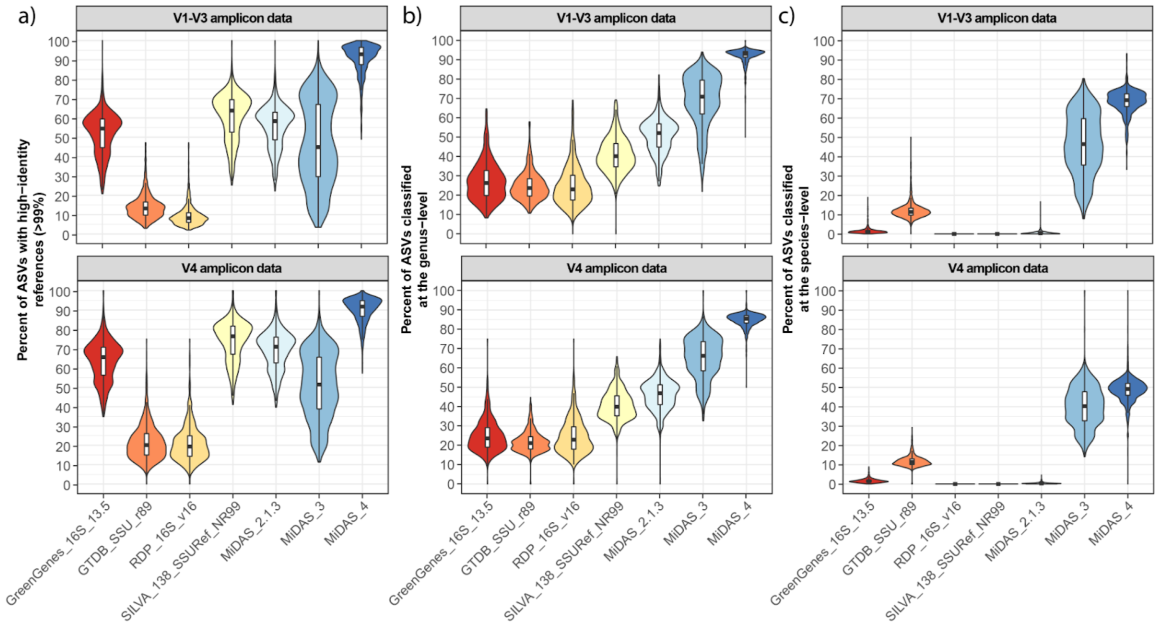


**Figure S2: Database evaluation based on amplicon data from this study.** V1-V3 and V4 ASVs were obtained from 1279 and 1278 samples, respectively, and filtered based on their relative abundance (≥0.01%) before the analyses. The percentage of the microbial community represented by the remaining ASVs after the filtering was 92.69% ± 3.22% and 96.4% ± 2.27% across samples for V1-V3 and V4 amplicon data, respectively. a) Percent of ASVs that have high-identity (≥99%) hits in MiDAS 4 and commonly applied universal reference databases. b) and c) Percent of ASVs that received b) genus- or c) species-level classification using sintax with the MiDAS 4 and commonly applied universal reference databases. Outliers have been removed from the boxplots.

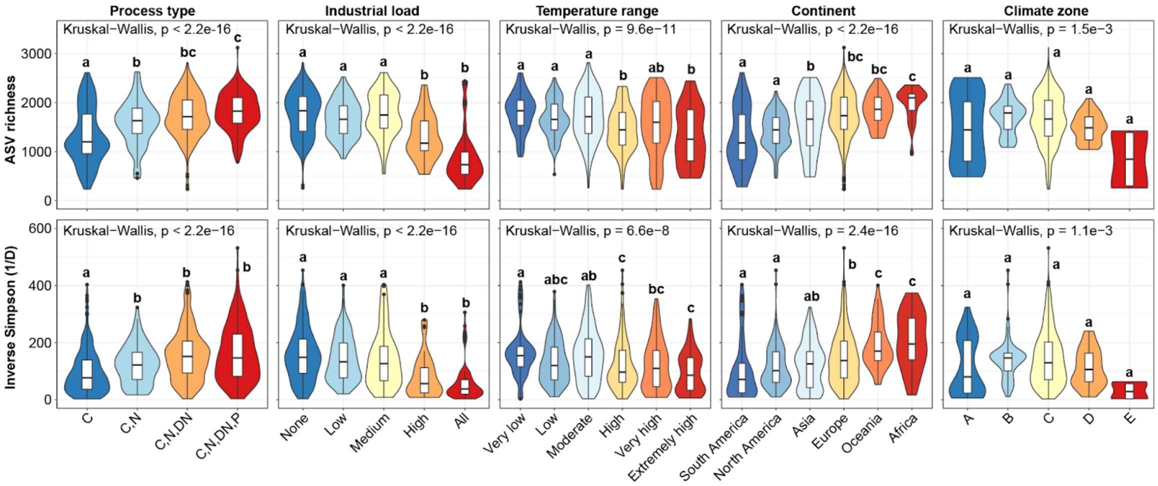


**Figure S3: Effect of process parameters and geography on alpha-diversity.** The non-parametric Kruskal-Wallis test was used to determine the statistical support for differences between the means of the groupings in the V1-V3 amplicon data set. A post-hoc Dunn's test (Bonferroni correction, α=0.01) was used for pairwise comparison of individual groups and the results are shown with compact letter display. Temperature range: very low = <10°C, low = 10-15°C, moderate = 15-20°C, high = 20-25°C, very high = 25-30°C, extremely high = >30°C . Industrial load: none = 0%, very low = 0-10%, low = 10-30%, medium= 30-50%, high 50-100%, all = 100%. Köppen climate classification groups: A = Tropical/megathermal climates, B = Dry (desert and semi-arid) climates, C = Temperate/mesothermal climates, D = Continental/microthermal climates, E = Polar climates. Outliers are not shown for the boxplots.

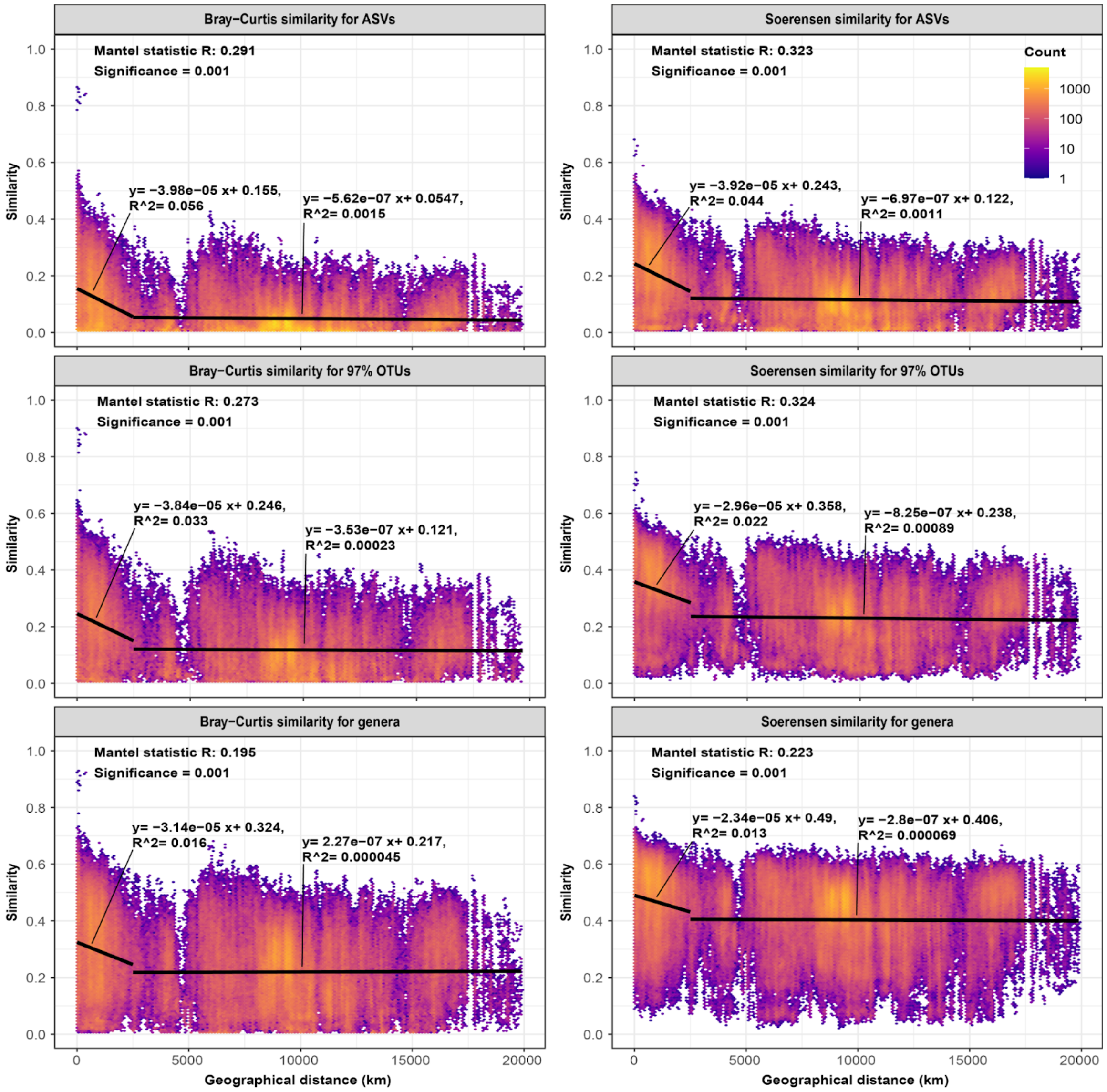


**Figure S4: Distance decay rates based on V1-V3 amplicon data.** a) DDR calculated based on Bray-Curtis (weighted) and Soerensen (unweighted) distances for ASVs, OTUs clustered at 97% sequence identity, and for genus-level classifications of ASVs. Mantel statistics for the DDR is provided for each plot. Each plot also contains the equations for two linear regressions. The first represent distances below 2,500 km and the second distances above 2,500 km.

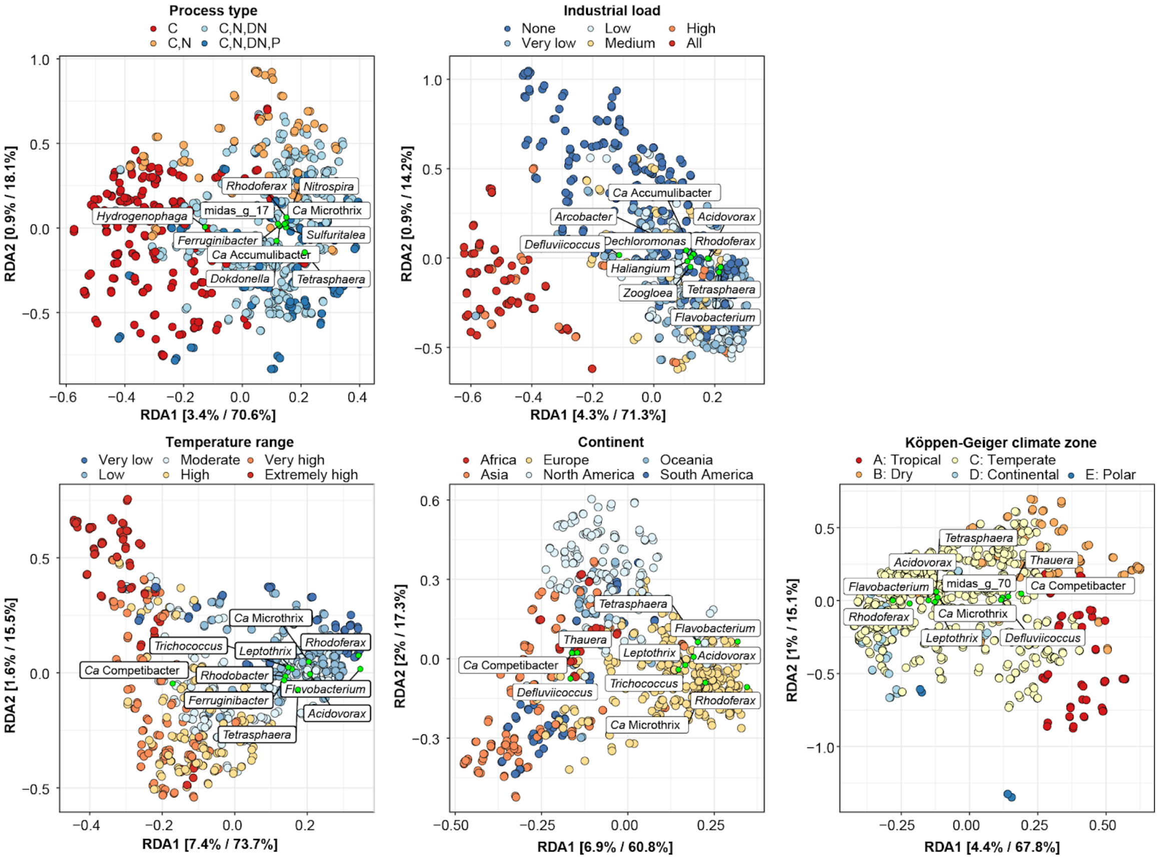

**Figure S5: Redundancy analyses (RDA) based on V1-V3 amplicon data classified at the genus-level constrained to each of the environmental or geographical parameters separately.** The data has been transformed initially by applying the Hellinger transformation. The relative contribution (eigenvalue) of each axis to the total inertia in the data, as well as to the constrained space only, respectively, are indicated in percent at the axis titles. Samples are colored based on metadata. The coordinates (green points) and names of the 10 most influential genera in relation to RDA1 is shown for each parameter. A comprehensive list with RDA scores for all genera can be found in **Supplementary Data S3**.

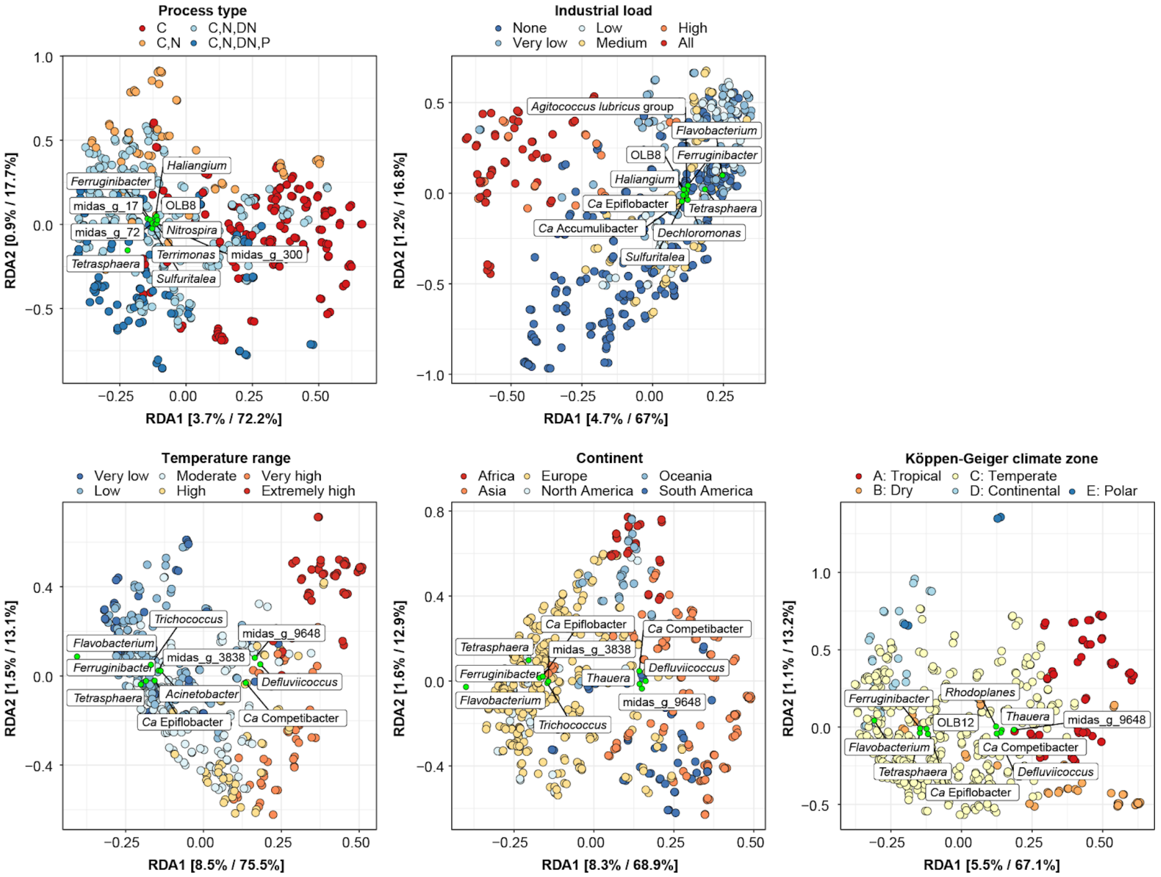


**Figure S6: Redundancy analyses (RDA) based on V4 amplicon data classified at the genus-level constrained to each of the environmental or geographical parameters separately.** The data has been transformed initially by applying the Hellinger transformation. The relative contribution (eigenvalue) of each axis to the total inertia in the data as well as to the constrained space only, respectively, are indicated in percent at the axis titles. Samples are colored based on metadata. The coordinates (green points) and names of the 10 most influential genera in relation to RDA1 is shown for each parameter. A comprehensive list with RDA scores for all genera can be found in **Supplementary Data S3**.

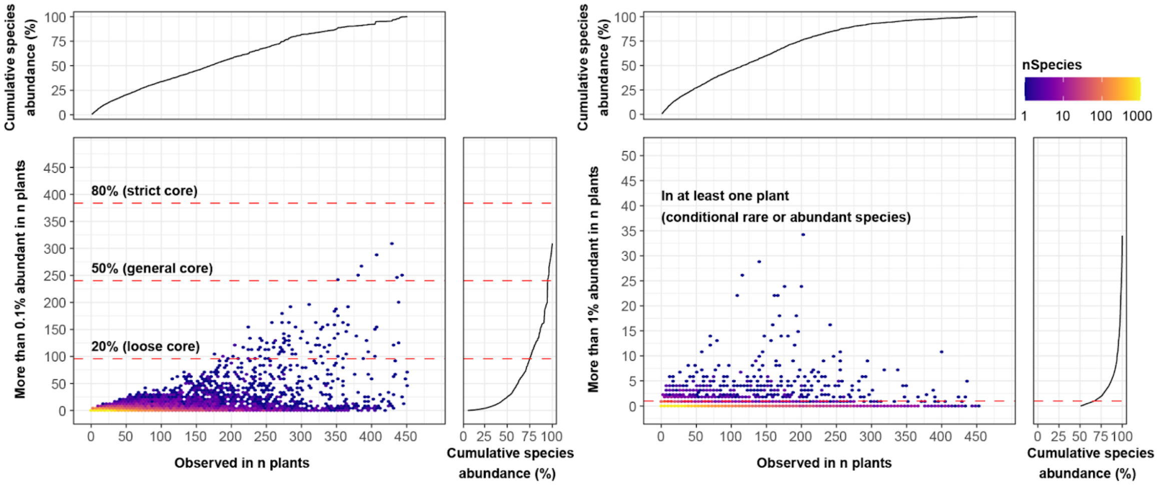


**Figure S7: Identification of core and conditionally rare or abundant species.** a) Identification of strict, general, and loose core species based on how often a given species was observed at a relative abundance above 0.1% in WWTPs. b) Identification of conditionally rare or abundant (CRAT) species based on whether a given species was observed at a relative abundance above 1% in at least one WWTP. The cumulative species abundance is based on all ASVs classified at the species-level. All core species have been removed before identification of the CRAT species.

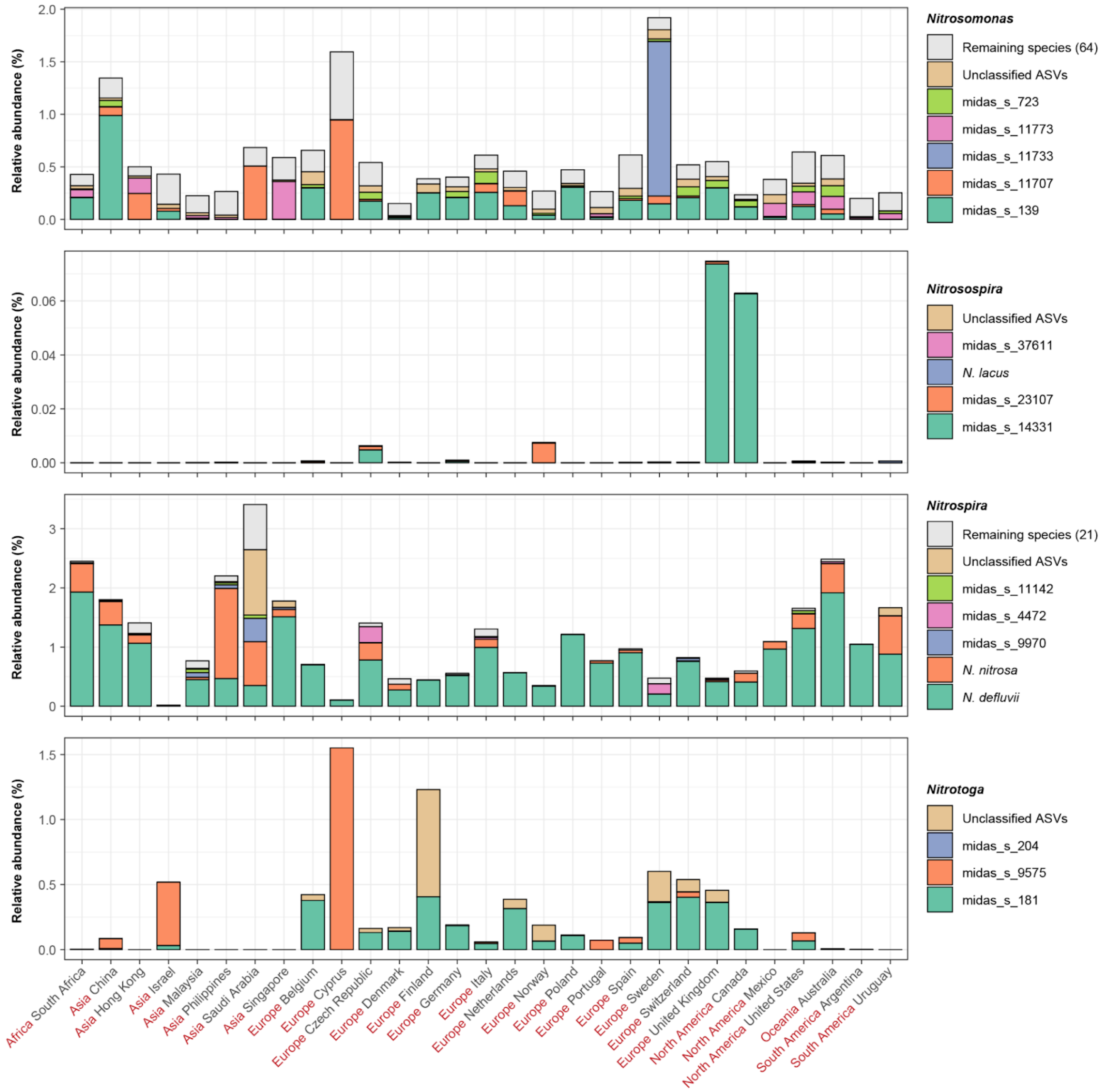


**Figure S8: Global species-level diversity of nitrifiers.** The percent relative abundance represents the mean abundance for each country taking into account only WWTPs with nitrification (C,N; C,N,DN, C,N,DN,P). The species are sorted based on their mean global abundance with the most abundant species at the bottom.

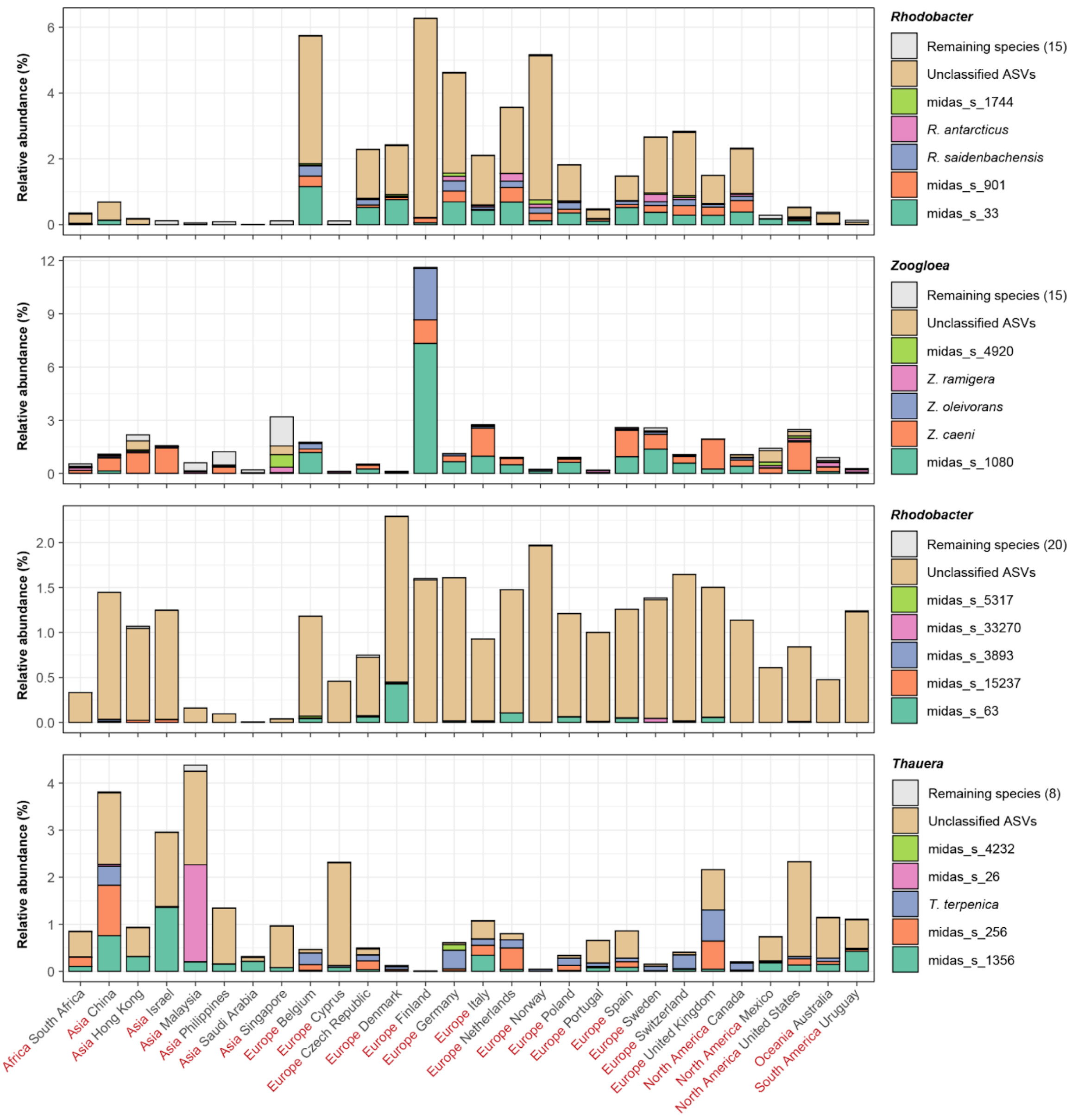


**Figure S9: Global species-level diversity of top four denitrifiers.** The percent relative abundance represents the mean abundance for each country taking into account only WWTPs with denitrification (C,N,DN, C,N,DN,P). The species are sorted based on their mean global abundance with the most abundant species at the bottom.

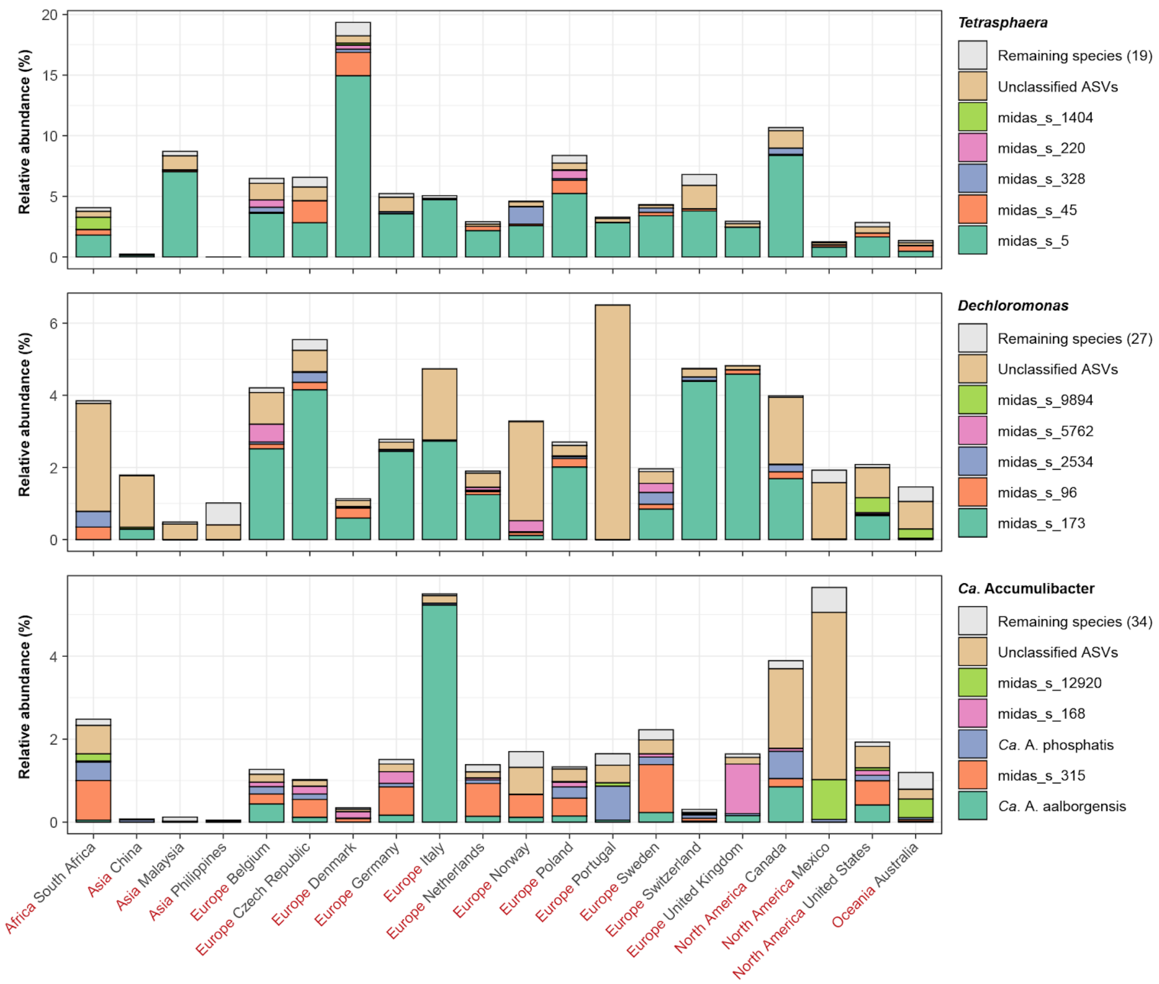


**Figure S10: Global species-level diversity of PAOs.** The percent relative abundance represents the mean abundance for each country taking into account only WWTPs with enhanced phosphorus accumulation (C,N,DN,P). The species are sorted based on their mean global abundance with the most abundant species at the bottom.

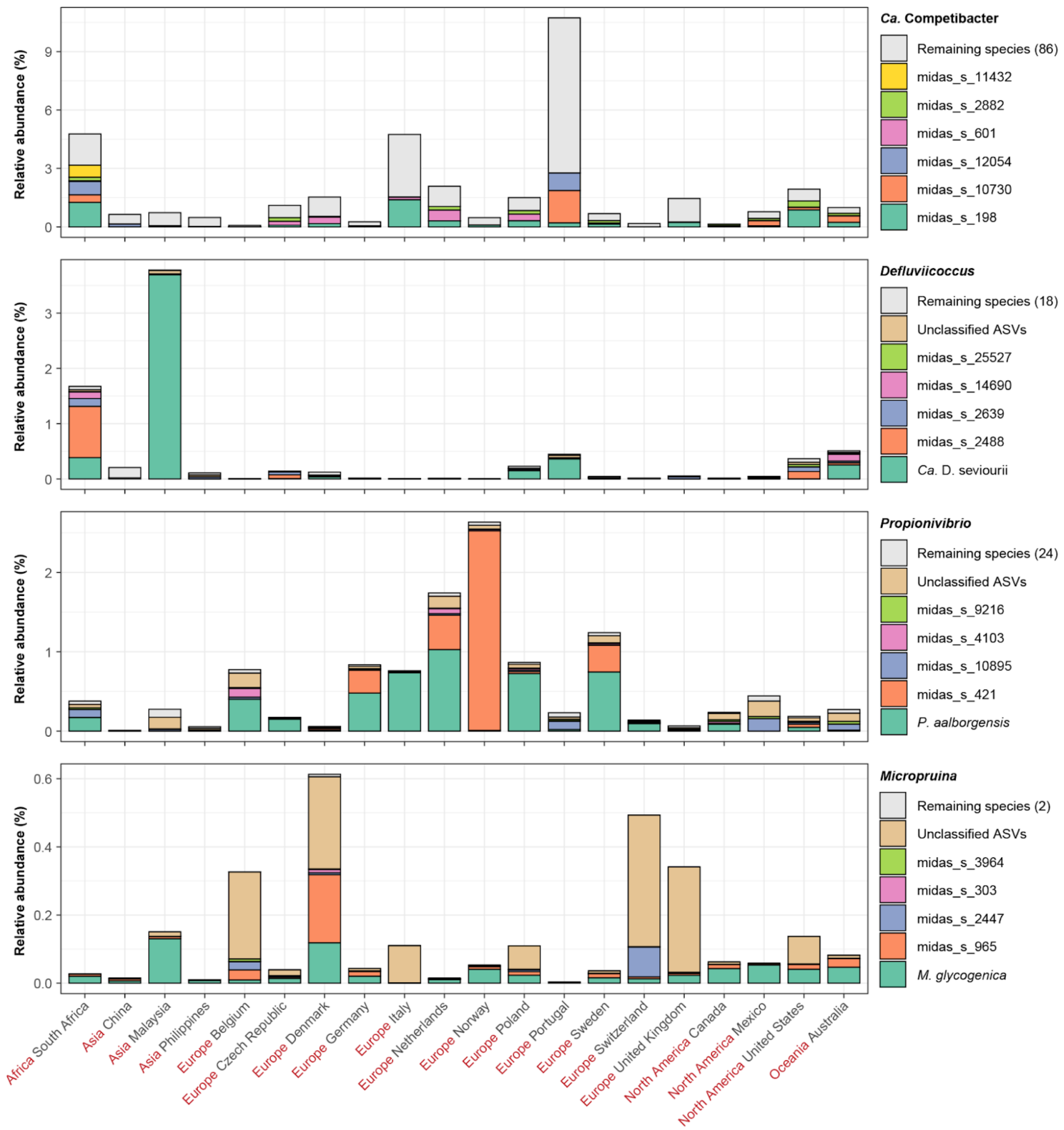


**Figure S11: Global species-level diversity of GAOs.** The percent relative abundance represents the mean abundance for each country taking into account only WWTPs with enhanced phosphorus removal (C,N,DN,P). The species are sorted based on their mean global abundance with the most abundant species at the bottom.

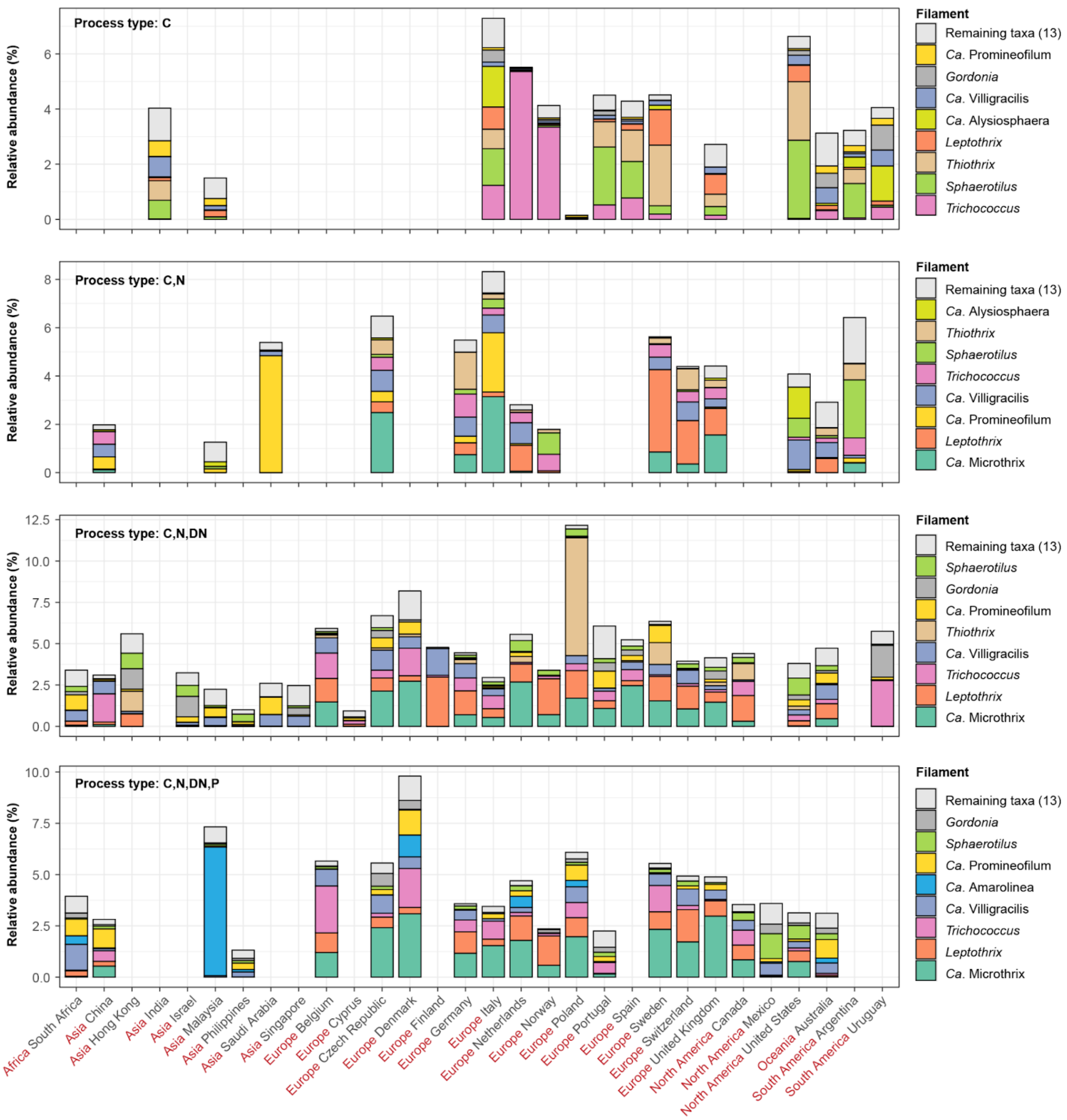


**Figure S12: Global diversity of known filamentous organisms in different process types.** The percent relative abundance represents the mean abundance for each country and each process type. The eight most abundant filamentous organisms based on mean relative abundance across the countries and process types are shown. The filaments are sorted based on their mean global abundance with the most abundant taxa at the bottom. The remaining taxa comprise *Haliscomenobacter*, *Defluviicoccus seviorii*, *Sarcinithrix*, *Ca.* Amarolinea, *Kouleothrix*, *Ca.* Alysiosphaera, *Nocardioides*, midas_g_1668, *Anaerolinea*, *Tetrashaera* midas_s_328, midas_g_105, midas_g_2111, midas_g_344, *Skermania*, *Ca*. Nostocoida, *Neomegalonema*, and *Beggiatoa* (not all were detected).

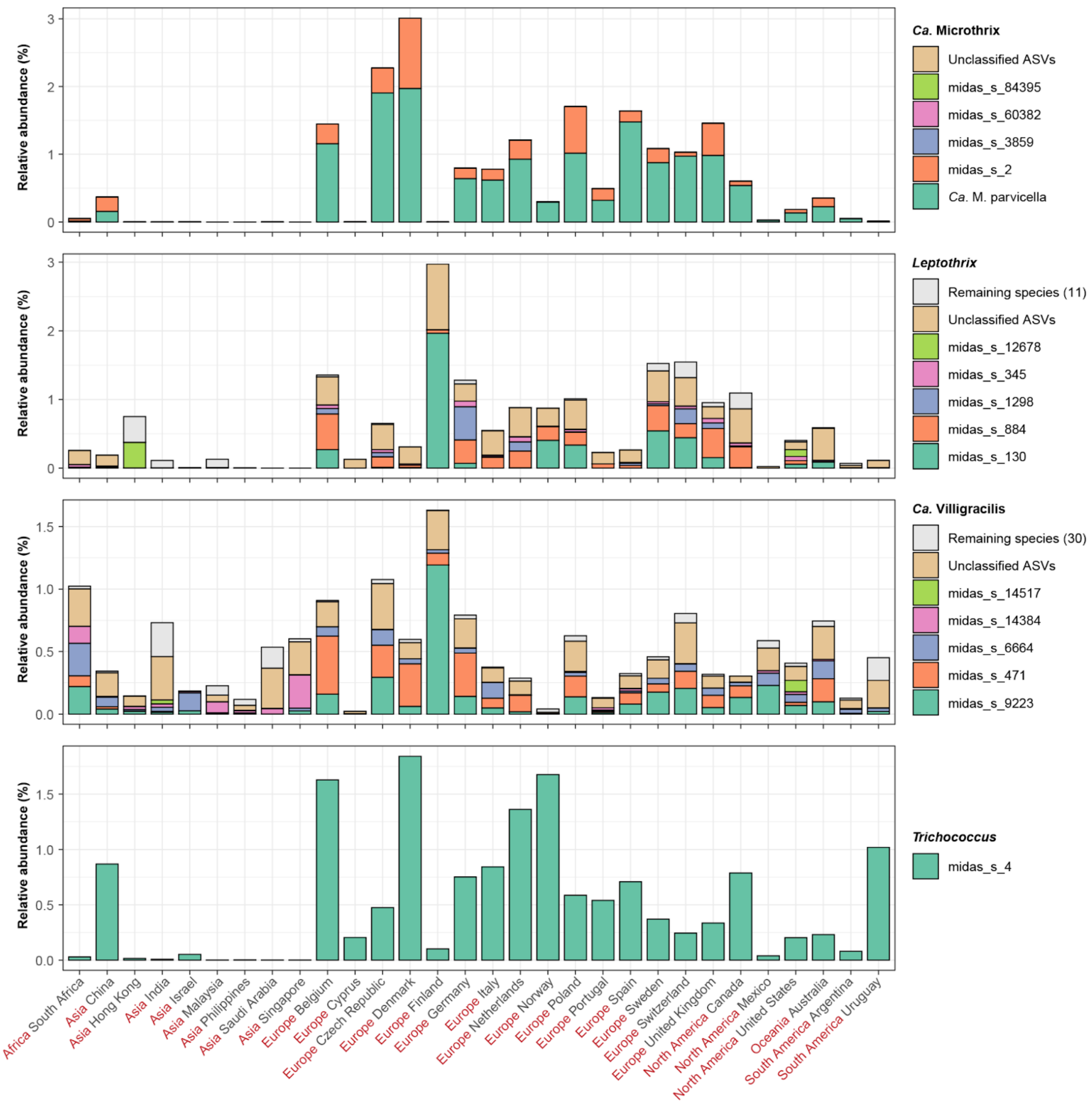


**Figure S13: Global species-level diversity of top four filamentous genera.** The percent relative abundance represents the mean abundance for each country. The species are sorted based on their mean global abundance with the most abundant species at the bottom.

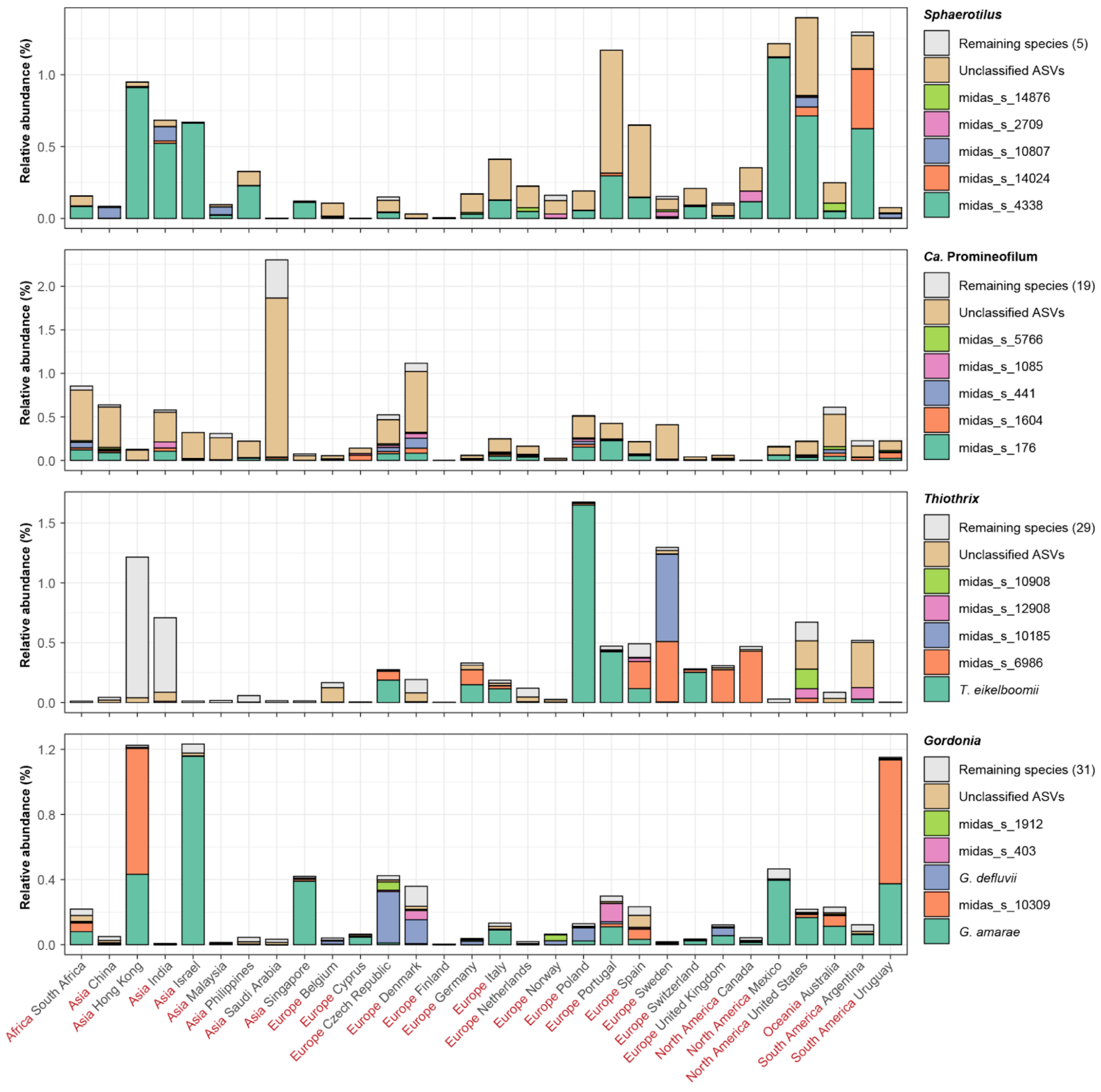


**Figure S14: Global species-level diversity of filamentous genera (top 5-8).** The percent relative abundance represents the mean abundance for each country. The species are sorted based on their mean global abundance with the most abundant species at the bottom.
